# A single residue determines regulation of TRPV1 by phosphoinositides

**DOI:** 10.1101/2025.10.15.682641

**Authors:** Benjamin He, Prateeti Varanasi, Nia M. Barkum, Rose Hudson, Eric N. Senning

**Affiliations:** Department of Neuroscience, University of Texas at Austin

## Abstract

The ion channel TRPV1 is expressed in the peripheral nervous system where it mediates heat sensation and pain signaling. How lipids contribute to TRPV1 regulation remains an open topic. Although an inhibitory binding site for phosphoinositides was identified at the vanilloid site in TRPV1 structures, a mechanism for phosphatidylinositol-2-bisphosphate (PI(4,5)P_2_) to both inhibit and potentiate TRPV1 activity is lacking. This gap in knowledge led us to examine structural and sequence overlap between TRPV1 and the PI(4,5)P_2_ binding site of TRPV5. In a space located adjacent to the turn from S6 to TRP helices, we identify a second, possible PI(4,5)P_2_ binding site in TRPV1, which we term the “front porch” site and includes residue H410. On the one hand, we obtained electrophysiological evidence with the TRPV1-H410R mutant that PI(4,5)P_2_ associates with TRPV1 at the front porch site through an electrostatic interaction to potentiate channel activity. On the other hand, the charge reversal in H410D conferred an increase in channel activity, which was not expected if the electrostatic interaction with PI(4,5)P_2_ is lost and channel function is crippled. Importantly, capsaicin activated currents from H410D experience significant current run-up, consistent with the slow displacement by capsaicin of a tighter-bound lipid in the vanilloid site, which structures suggest is phosphoinositide (PI). Lastly, we tested TRPV1-H410D sensitivity to PI(4,5)P_2_ manipulations by dephosphorylating the membrane pool of PI(4,5)P_2_ to PI with the Pseudojanin rapamycin-phosphatase (PJ) system. Depletion of PI(4,5)P_2_ with the PJ-system in wild type channels induces 1) a robust increase in baseline activity as channels are released from inhibition and 2) loss of current potentiation, driven by PI(4,5)P_2_ dissociation from the front porch, both of which were absent in TRPV1-H410D. Based on these findings we hypothesize that plasma membrane TRPV1-H410D maintains PI rather than PI(4,5)P_2_ in its vanilloid site and does not interact with PI(4,5)P_2_ in the front porch site. We integrated our discoveries that H410 interacts with PI(4,5)P_2_ at two distinct sites in a mutually exclusive manner and present a cohesive model for phosphoinositide inhibition and potentiation in TRPV1, addressing a maturation process for TRPV1 that relies on H410 to coordinate the exchange of PI for PI(4,5)P_2_.

**Statement of Significance:** A lingering question that hangs over TRPV1 research is how the lipid PI(4,5)P_2_ regulates the function of this channel. We devised a study based on structural homology between TRPV5 and TRPV1 to identify critical contact points between the lipid and a putative PI(4,5)P_2_ binding site in TRPV1, which we probed using mutagenesis experiments. The results we obtained lay the groundwork for how a two-site binding mechanism of PI(4,5)P_2_ couples to TRPV1 gating and address important questions about the channel’s maturation and association with phosphoinositides as it is transferred through membrane compartments with different phosphoinositide compositions. We conclude that TRPV1 expressed and purified from different compartments may preserve the phosphoinositide character of that compartment (PI in the endoplasmic reticulum, PI(4,5)P_2_ in the plasma membrane) and bestow on structures a state that is not necessarily relevant to the biological function of the channel in the plasma membrane.

## Introduction

The transient receptor potential vanilloid subtype I (TRPV1) channel is a cation channel expressed in sensory ganglia and known for its role in temperature sensation and pain transduction[1, 2]. TRPV1 has an exceptional activation profile and can be described as polymodal for the array of stimuli that modify or induce gating, including heat, oxidation, acidity, and exogenous vanilloids[3]. The range of activators also includes lipids, which, given their diverse make-up, have a spectrum of effects on TRPV1[4]. Although some of the minor lipids such as products of arachidonic acid metabolism and lysophosphatidic acid are shown to increase TRPV1 function[5, 6], the role of major lipids in the plasma membrane such as phosphatidylinositols remains at the focus of numerous research groups with competing mechanistic models for how these lipids interact with the channel to affect gating[7].

Early studies examining the role of GPCR and TrkA receptor signaling on TRPV1 function implicated phosphatidylinositol-2-bisphosphate (PI(4,5)P_2_) as an inhibitor of the channel. Focus on PI(4,5)P_2_ was further enhanced in this study by hydrolyzing PI(4,5)P_2_ directly with recombinant PI-PLC or sequestering the lipid with a PI(4,5)P_2_ antibody in excised patches [8]. Contradictory results were obtained in electrophysiology studies using direct application of phosphoinositide lipids onto the inner leaflet of excised patches with expressed TRPV1. First, plasma membrane sequestration of PI(4,5)P_2_ reduced TRPV1 currents and addition of short-chained dic8-PI(4,5)P_2_ to the inner leaflet restored the currents [9, 10]. Moreover, current restoration in the membrane patch was contingent on the type of polyphosphoinositide that was tested and which leaflet of the membrane was exposed to the short-chain lipid. Cellular manipulation of phosphoinositides evoked similar changes to TRPV1 current as would be expected if PI(4,5)P_2_ potentiates activity. Rapamycin-based systems that recruited the phosphoinositide phosphatases to the plasma membrane that convert PI(4,5)P_2_ to PI(4)P and phosphoinositide (PI) induced a reduction in TRPV1 currents[11, 12].

Subsequent electrophysiology with reconstituted TRPV1 in vesicles recapitulated that different phosphoinositides could evoke a range of effects on TRPV1 current. However, the capsaicin dose-response profiles obtained with TRPV1 in reconstituted membranes containing PI(4,5)P_2_ showed decreased sensitivity to capsaicin relative to phosphoinositide-free membranes, implicating the lipid as inhibitory [13]. Structural studies of TRPV1 also present a PI binding site in closed channel structures that would otherwise overlap with the vanilloid site where capsaicin and RTX agonists bind in open channel structures[14-16]. A recent structure obtained with brominated PI(4,5)P_2_ in TRPV1 confirms that this lipid is accommodated in the same position as the PI[17]. Structural evidence is, therefore, supportive of phosphoinositides in the vanilloid site as inhibitory since PI and PI(4,5)P_2_ bound structures have been solved in states with pores classified as non-permeant. To complement this, numerous TRPV1 structures have been solved with the PI displaced by capsaicin or RTX in the vanilloid binding site with a pore that is dilated and, potentially, in a permeant state[16, 18, 19].

The broad scope of lipid research on TRPV1 sheds light on three aspects of phosphoinositide regulation over TRPV1 that need to be reconciled: 1. Low activity of TRPV1 can be increased by releasing channels from inhibition with PI(4,5)P_2_ removal from the plasma membrane. 2. TRPV1 currents activated by near-saturating concentrations of capsaicin diminish when PI(4,5)P_2_ is depleted from the plasma membrane; 3. Structural studies of TRPV1 show densities attributable to PI and PI(4,5)P_2_ overlaping with the vanilloid binding site that implicate a competitive activity for the phosphoinositides. Yazici and colleagues attempted to discern how PI(4,5)P_2_ and PI interact with the channel to reconcile these contradictory findings and hypothesized that two binding sites exist for the lipids[20]. Their research relied on molecular dynamics (MD) simulations of PI and PI(4,5)P_2_ in TRPV1 structures, resolving two possible binding sites: One site was identified as inhibitory, being preferred by PI and overlapping with the vanilloid binding site; a second site was preferred by PI(4,5)P_2_, further removed from the inhibitory PI binding site, and designated as activating. Although this study hypothesized the existence of two binding sites, mechanistic descriptions of their functional attributes that lead to channel gating remain incomplete.

Here we present new results that follow from our recent MD study done with TRPV5 and PI(4,5)P_2_ and analysis of structural alignments between TRPV5 with TRPV1[21]. TRPV5 is a member of the vanilloid subfamily of TRP channels having 30.6% amino acid sequence identity with TRPV1 (See methods). PI(4,5)P_2_ activates TRPV5, and a well-resolved structure shows the location of a dic8-PI(4,5)P_2_ molecule bound to the channel (**Figure 1A,B**; PDB: 6DMU, cyan)[22]. Our previous MD study examined the pH dependence of PI(4,5)P_2_ association with TRPV5 through the Milestoning technique and characterized the relevant amino acid side-chain interactions with the lipid[21]. Along with several residue-lipid interactions already noted in the TRPV5 structure, our research highlighted residues that interact with the PI(4,5)P_2_ as it approaches the binding pocket and pinpointed the side-chain of R305 as a critical, pH dependent contact between the 5’ phosphate of PI(4,5)P_2_ and channel. Sequence and structural alignment with TRPV5 identified H410 in TRPV1 as a match to R305, and we show through functional studies that H410 is a determinant of phosphoinositide regulation over TRPV1 function. Our study brings the lipid regulation of TRPV1 into sharper focus with the discovery of the mutant TRPV1-H410D that appears to have lost all PI(4,5)P_2_ dependent regulation while retaining capsaicin dependent activation. Results from experiments with TRPV1-H410 mutants allowed us to construct a two-site phosphoinositide binding model with the side chain of H410 forming instrumental contacts with each site to facilitate both inhibition and potentiation in a mutually exclusive manner. We then apply this model to explain TRPV1 maturation in cells.

**Figure 1.**
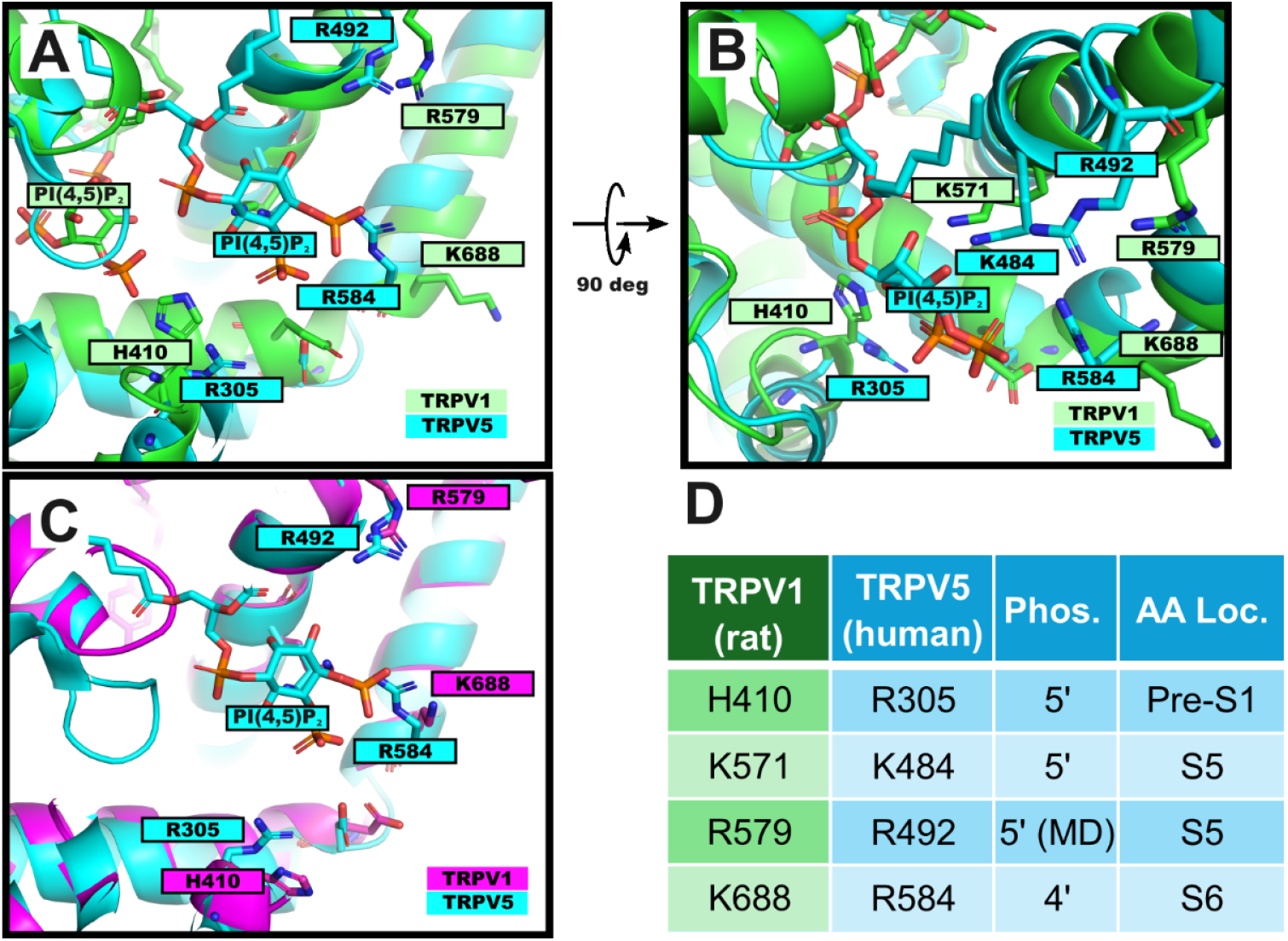
Structural similarity of a PI(4,5)P_2_ binding site in TRPV5 and TRPV1. (A) Side perspective of the PI(4,5)P_2_ binding pocket in TRPV5 (PDB:6dmu, blue) aligned with a dic8-PI(4,5)P_2_, closed state structure of TRPV1 (PDB: 8u30, green). Labeled side chains of TRPV5 residues are selected for their extensive interactions with PI(4,5)P_2_ as defined in Fathiyadeh et al (2021). Analogous side chains of TRPV1 are shown in green (B) Top perspective of PI(4,5)P_2_ binding site shown in panel (A). (C) TRPV5 PI(4,5)P_2_ binding pocket shown in alignment with an RTX/DkTx bound, open-state TRPV1 structure (PDB:7l2m, magenta). (D) Table of residues identified in panels A-C with listing of interacting phosphate group in PI(4,5)P_2_ and topological location in channel.

## Materials and Methods

### Sequence data, genomic variants and structural analysis

Human genomic variant data were obtained from the gnomad database (https://gnomad.broadinstitute.org/). Sequence identity between human TRPV1 (Uniprot ID: Q8NER1) and human TRPV5 (Uniprot ID: Q9NQA5) was calculated as the percent identity matrix through an alignment of the protein sequence with the Uniprot alignment tool (https://www.uniprot.org/align). Structure files were aligned (residues 300-600 in one TRPV5 [PDB: 6dmu] subunit were aligned with residues 400-700 in one TRPV1 [PDB: 8u30;7l2m] subunit) and rendered for publication with PyMOL (Schrödinger, NY). Gene sequence data was acquired through the NCBI.NLM.NIH.gov and aligned in Seaview for publication[23].

### Cell culture

HEK293T/17 (ATCC: CRL-11268) cells were incubated in Dulbecco’s Modified Eagle Medium, supplemented with 10% fetal bovine serum, 50 μg <u>streptomycin</u>, and 50 units/mL penicillin, at 37°C and 5% CO2. Cells were passaged onto poly-L-lysine treated 25 mm or 10 mm coverslips. Cells were allowed at least 2h to settle onto the slips before being transfected using <u>Lipofectamine</u> 2000 (Life Technologies) as described in the manufacturer’s instructions. Cells were transfected with pcDNA3.1 containing TRPV1 wild type and mutants. Following transfection, Ca^2+^ imaging experiments were carried out after 24h, and electrophysiology experiments were completed 24-48h after transfection. For electrophysiology experiments co-transfection of the TRPV1 construct of interest was done alongside a GFP expressing plasmid (JSM-164 NW10 3C-GFP-HS)[24].

### Molecular biology

In vivo assembly (IVA) methods of site-directed mutagenesis were done as previously described[25]. For all in-house mutagenesis, our wild-type TRPV1.pcDNA3 was used as a template with (IVA) overlapping primers that contain the desired mutation (see primers for mutagenesis spreadsheet). The full sequence of each completed construct was confirmed using Whole Plasmid Sequencing (Plasmidsaurus, CA).

### Electrophysiology

#### Capsaicin dose response experiments at nominal and high pH

Currents were recorded at room temperature in symmetric divalent-free solutions (130 mM NaCl, 3 mM HEPES, 0.2 mM EDTA, pH 7.2) using fire polished borosilicate glass pipettes with filament (outer diameter 1.5 mm, inner diameter 0.86 mm; Sutter, MA). Using a micro forge, for excised patch experiments, pipettes were heat polished to a resistance of 3.5-7 MΩ and for whole cell experiments, pipettes were heat polished to a resistance of 2-4 MΩ. Cells were transfected with channel constructs and replated after 18-24 hours on 12 mm coverslips and placed in divalent-free solution in the chamber. 1, 3, 5, and 10 µM capsaicin solutions were prepared from a 5.3 mM capsaicin (Sigma Aldrich, MO) stock in ethanol. A dilution of the 5.3 mM stock down to 0.5 mM capsaicin in ethanol stock solutions was used for preparing 0.03 µM, 0.1 µM, and 0.3 µM capsaicin solutions in divalent free recording solution. Excised, inside-out patches were positioned at the opening of a capillary glass tube in a “sewer pipe” configuration with solution tubes braced to a rod controlled by an RSC-200 rapid solution changer (BioLogic, France). Each successive perfusion tube was briefly primed by applying pressure through the syringe reservoir before the tube was positioned in front of a patch. Capsaicin solutions (0.03, 0.1, 0.3, 1, 5 and 10 µM) were perfused using open, gravity-drive reservoirs for 40-120s each. pH dependent experiments involving WT TRPV1 and each variant were conducted using divalent-free HEPES buffer as described above with pH adjusted as indicated in experiments (nominal pH: 7.2; high pH: 8.0). Currents were measured with an Axopatch-200A e-phys amplifier system and PatchMaster software (HEKA). Command voltage sweeps were done at intervals of 1 second stepping the voltage from resting potential (0mV) to −80 mV (300 ms), then up to +80 mV (300 ms) and back to resting potential. Currents were filtered at 5 kHz and recorded at 20kHz using an Axopatch 200A amplifier (Axon Instruments, Inc.) and. Data was analyzed in PatchMaster and IGOR (Wavemetrics, OR). Individual patch-clamp recordings with a complete set of stable current measurements in all capsaicin solutions were normalized to the maximal current with 5 µM or 10 μM (H410D) capsaicin. Normalized data was fitted to the Hill equation to arrive at the EC_50_ parameter: *I* = *I*_max_ ([ligand]^*n*^/(EC_50_^*n*^ + [ligand]^*n*^)), where n was permitted to float. Statistical significance in Figures 2 and 3 was determined by two sample, one-tailed Student’s t-tests.

**Figure 2.**
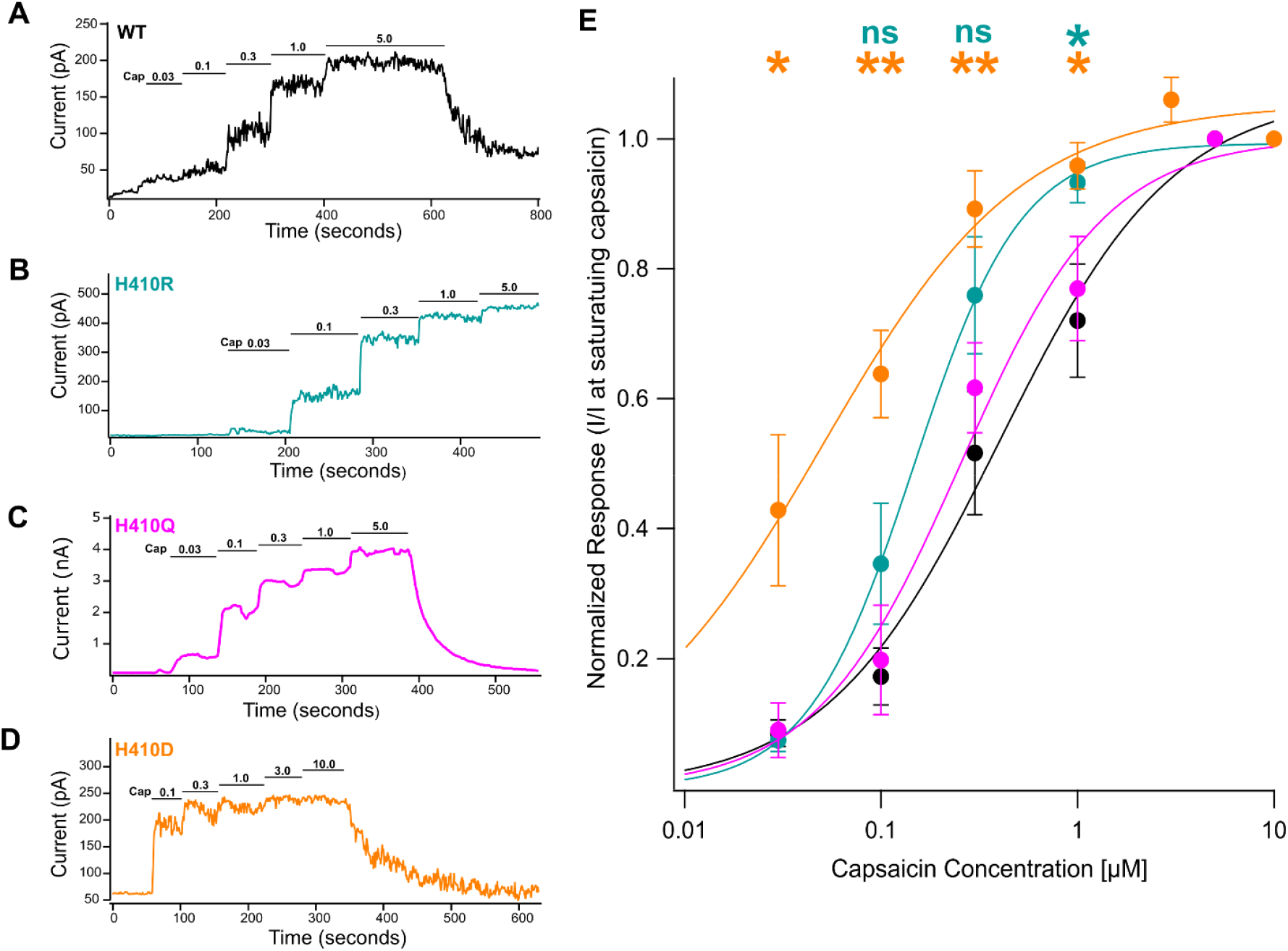
Both TRPV1-H410R and TRPV1-H410D have increased sensitivity to capsaicin relative to wild type. (A-D) Representative traces of capsaicin dose response relations in TRPV1-WT and TRPV1-H410 mutants (TRPV1-H410R, TRPV1-H410Q, and TRPV1-H410D respectively). Solution exchanges with different concentrations of capsaicin given in μM above traces. (E) Summary data of capsaicin dose response experiments shown as log-linear plots and fitted with a Hill function to acquire EC_50_ values (in μM: TRPV1-WT = 0.40286, TRPV1-H410R = 0.14698, TRPV1-H410Q = 0.25515, TRPV1-H410D = 0.050083). The TRPV1-H410D mutant average response is significantly different from wild type TRPV1 at four capsaicin concentrations, 0.03, 0.1, 0.3, and 1.0 μM (p = 0.015796, 0.000534, 0.006656, 0.020141. Student’s two sample, one-tailed t-test). The TRPV1-H410R mutant average response is significantly different from wild type TRPV1 at 1.0 μM capsaicin (“*” at 1 μM: p = 0.0382471; ‘ns’: not significant at 0.1 μM: p = 0.056235, at 0.3 μM: p = 0.055890). Student’s two sample, one-tailed t-test. Error bars are S.E.M.

**Figure 3.**
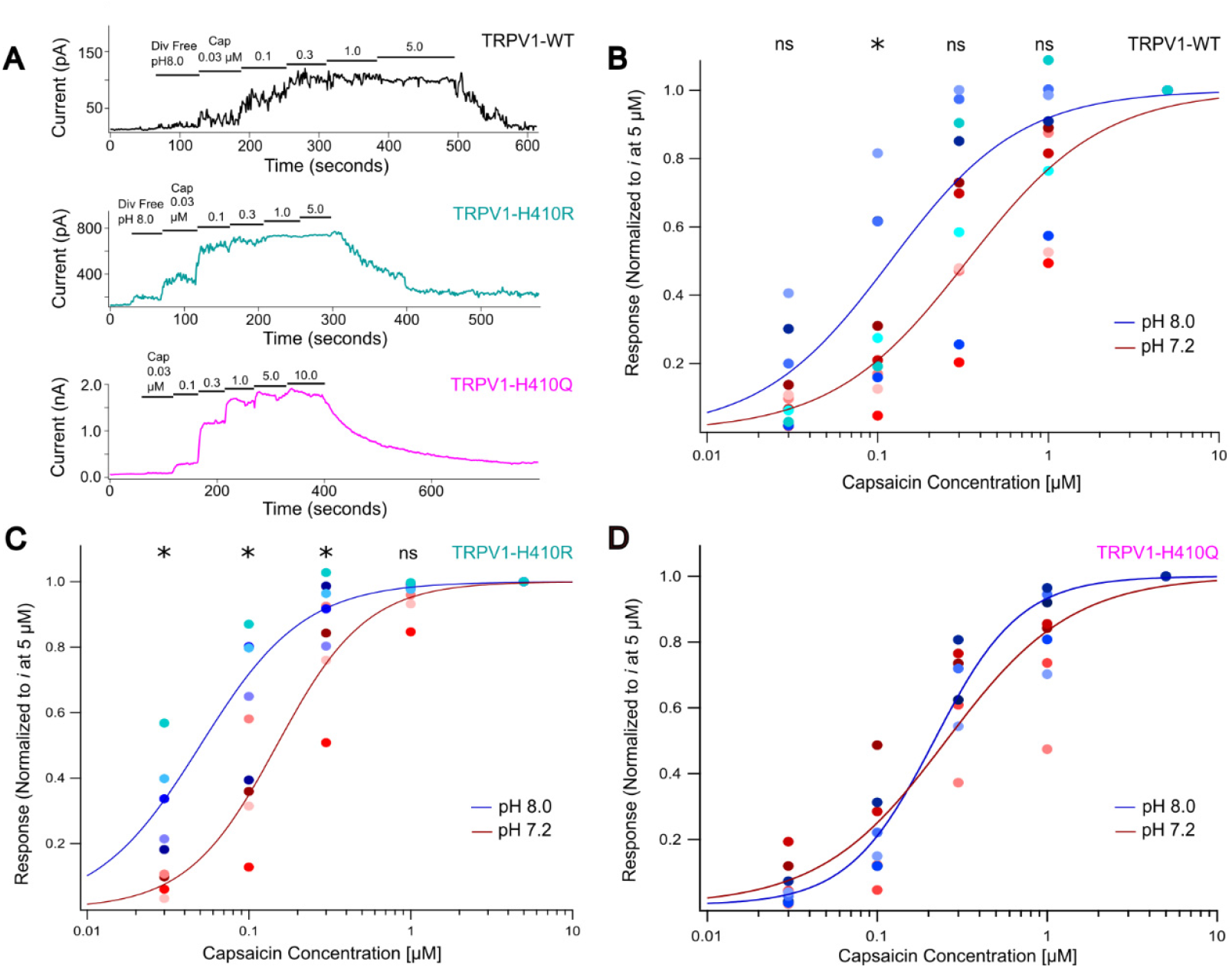
Current sensitivity to inner leaflet pH is lost in TRPV1-H410Q. (A) Representative capsaicin dose responses traces collected using pH 8.0 solutions shown. Some experiments began with a brief application of pH 8.0 divalent free solution free of capsaicin before applying the capsaicin solution set. B) Summary data of capsaicin dose response behavior for TRPV1-WT is shown at both pH 7.2, represented by shades of red, and pH 8.0, represented by shades of blue. Data was represented with log-linear plots and fitted by a Hill equation to derive EC_50_ values. In pH 7.2 and pH 8.0 solutions, the EC_50_ of TRPV1-WT occurred at 0.335 µM capsaicin and 0.119 µM capsaicin, respectively. The average response to capsaicin across the two pH levels was significantly different at one capsaicin concentration, 0.1 µM (‘*’: statistically significant at p = 0.0317; ‘ns’: not significant at 0.03 µM p = 0.144; at 0.3 µM p = 0.0750; at 1 µM p = 0.0911. Student’s two sample, one-tailed t-test). C) Summary capsaicin dose response data for TRPV1-H410R is shown using the same color scheme and analysis. In pH 7.2 and pH 8.0 solutions, the EC_50_ of TRPV1-H410R occurred at 0.149 µM capsaicin and 0.0501 µM capsaicin, respectively. The average response between the two pH levels was significantly different at three capsaicin concentrations: 0.03 µM, 0.1 µM, and 0.3 µM (‘*’: statistically significant at p = 0.00645, p = 0.0128, p = 0.0429; ‘ns’: not significant at 1 µM p = 0.0501. Student’s two sample, one-tailed t-test). D) Summary capsaicin dose response data for TRPV1-H410Q is shown using the same color scheme and analysis. In pH 7.2 and pH 8.0 solutions, the EC_50_ of TRPV1-H410Q occurred at 0.255 µM capsaicin and 0.216 µM capsaicin, respectively. The average response between the two pH levels was not significantly different at any capsaicin concentration (at 0.03 µM p = 0.0896; at 0.1 µM p = 0.439; at 0.3 µM p = 0.295; at 1 µM p = 0.294. Student’s two sample, one-tailed t-test).

#### Poly-lysine and magnesium experiments

With fewer tubes used in the Biologic sewer pipe perfusion system, TRPV1 channel currents in excised patches from HEK293T cells were recorded in response to 1.0 µM capsaicin divalent free HEPES (preparation described above) with and without 50 mM MgCl_2_ added to the capsaicin solution. Poly-lysine inhibition experiments were done with and without 3 µg/mL poly-lysine in the base, capsaicin solution. The capsaicin solution was made as detailed above with 5.3 mM stock in divalent-free HEPES buffer. A 2 M MgCl_2_ stock was used to prepare a 50 mM MgCl_2_ in 1.0 µM capsaicin solution. Similarly, a 2.0 mg/mL stock of poly-lysine was used to prepare the 3.0 µg/mL poly-lysine in 1.0 µM capsaicin solution. Divalent-free pH 7.2, 1.0 µM capsaicin, and the MgCl_2_ with capsaicin solutions were perfused for 30-120s each with an open, gravity-driven reservoir via the sewer pipe system onto excised membrane patches. In some experiments, the order of stepping potentials was reversed but this did not alter the outward current used to measure TRPV1 activity. Individual patch-clamp recordings with a complete set of stable current measurements in all solutions were adjusted for leak and the ratios of current with magnesium and capsaicin over only capsaicin were calculated. Statistical significance was determined by two sample, two-tailed Student’s t-tests.

#### Capsaicin run-up experiments

Using the Biologic sewer pipe perfusion system as described above, channel currents in excised patches from HEK293T cells expressing wild type or mutant TRPV1 channels were recorded in response to extended application of high (3.0 μM) capsaicin solutions. 0.1 and 3.0 μM capsaicin solutions were prepared from a 5.3 mM capsaicin stock in ethanol. Capsaicin solutions were perfused at varying time frames with 0.1 µM perfusing for 30-60s, 3.0 µM perfused for 120-180s then another perfusion of 0.1 µM for 30-60s. After leak subtraction, the ratio of the final currents collected at 0.1 µM capsaicin occurring after compared to before the 3.0 µM capsaicin was examined for a run-up effect. Statistical significance of change between 0.1 µM capsaicin currents after and before 3.0 µM capsaicin was evaluated by comparing the ratio of before and after currents to 1.0 with a Student’s one-sample, two-tailed t-test.

#### Whole cell electrophysiology measurement of TRPV1 currents with rapamycin induction of Pseudojanin phosphatase system

HEK293T cells that were previously plated onto 10mm coverslips were transfected with the PJ system constructs mRFP-FKBP-Pseudojanin(PJ; Addgene 37999), Lyn_11_-FRB-CFP (Addgene; 38003) and the TRPV1 construct of interest at the recommended ratio of 5:3:3 (Lyn_11_:TRPV1:PJ) with Lipofectamine 2000 reagent according to manufacturer’s instructions[12]. After 24 hours, cells were transferred in divalent free HEPES solution (preparation described above) to an experimental chamber fitted with gravity fed perfusion for whole cell electrophysiology. Divalent free HEPES solution was used as intracellular pipette solution to maintain symmetrical solution conditions with the bath. Whole cell electrophysiology pipettes were heat polished to a resistance of 2-4 MΩ. Break-in to whole cell patch clamp configuration was done with >200MΩ seal resistance, and outward currents were measured with an Axon200A amplifier and the same voltage sweep protocol as described for the excised patch, capsaicin dose-response experiments.

Before testing the PI(4,5)P_2_ dependence of current in TRPV1 channel constructs with the PJ-rapamycin system, one set of current-voltage plots was collected to verify the presence of an outwardly rectifying current that is the hallmark of TRPV1 expression. Experimental capsaicin solution (3.0 μM) was prepared in divalent free HEPES from a 5.0 mM ethanol stock and this 3.0 μM working, capsaicin solution was used to prepare a 1 μM rapamycin and 3 μM capsaicin solution from a stock of 5.5 mM rapamycin dissolved in DMSO (Millipore, USA). After 50 seconds in divalent free solution to record baseline currents, cells were perfused with the 3.0 μM capsaicin solution for 70-100 seconds to activate channels at near saturating conditions. To stimulate PI(4,5)P_2_ depletion with PJ, we switched perfusion to the capsaicin/rapamycin solution for 150-200 seconds to ensure complete equilibration of the rapamycin concentration in the bath chamber. Complete washout of the capsaicin and rapamycin with divalent free HEPES solution proceeded at variable rates depending on gravity flow rate but also clustering of cells around patched cell in the experiment. Patched cells that exceeded >10 nA of current with the initial application of 3.0 μM capsaicin were rejected. Statistical significance was evaluated with Student’s two-sample, one-sided t-test.

### Calcium Imaging

#### TRPV1 mutant screening

Cells were plated on 25-mm coverslips before being transfected as previously described. After 24 hours, transfected cells were incubated for 30 min at 37 °C with Fluo-4 (AM; Thermofisher) at a concentration of 3 μM. The cells were then rinsed with Hepes buffered Ringer’s (HBR) solution (in mM, 140 NaCl, 4 KCl, 1.5 MgCl2, 5 D-glucose, 10 HEPES, and 1.8 CaCl2 and pH adjusted to 7.4 with NaOH) and allowed to rest for another 30 min in HBR at room temperature. The cells were imaged on a Nikon Eclipse Ti microscope using a 10 × objective. For each slip, a brightfield image, a fluo-4 image, and a 3-min fluo-4 fluorescence movie with exposures of 100-ms and 0.5-s intervals were captured. During the time sequence, HBR is initially perfused throughout the chamber. Perfusion is switched to 500 nM capsaicin in HBR at 30 s, and at the 2 min mark, 3 μM ionomycin is added to the chamber to equilibrate calcium between inside and outside the cell in order to obtain the maximal Fluo-4 response. The HBR and 500 nM capsaicin in HBR were perfused into the chamber using open, gravity-driven reservoirs, and 500 uL of 3 μM ionomycin was Pipetted into the chamber via micro-Pipette. The data obtained with these experiments were analyzed using Image J and Excel (Microsoft, WA).

#### Calcium imaging experiments with TRPV1 constructs and rapamycin induction of Pseudojanin phosphatase system

Cells were plated on 25 mm coverslips and transfected overnight with TRPV1 constructs and PJ-system as described in whole cell TRPV1 with PJ-rapamycin experiments. Fluo4 loading proceeded as previously described for calcium imaging with HBR solution and cells were mounted in an imaging chamber fitted with gravity fed perfusion. A 1.0 mM rapamycin solution was prepared in HBR from 5.5 mM stock in DMSO (Millipore, USA) and a 0.1 mM capsaicin solution was prepared in HBR from a 5.0 mM stock in ethanol. Imaging proceeded with one image of the CFP signal to capture cells expressing Lyn_11_-FRB-CFP, one image of the RFP signal to capture cells expressing mRFP-FKBP-Pseudojanin and a time series of the Fluo4 signal using a GFP excitation/emission cube. The image series of Ca^2+^ signal was captured at 2 Hz with EM gain set to 50 on the Ixon Life camera (Andor, UK). After 30 seconds in HBR to establish a baseline signal, the rapamycin solution was rapidly introduced to the chamber, reaching equilibrium within 20 seconds. At the 2 minute 30 second mark, perfusion switched to the capsaicin solution. At the 4 minute mark 5 μM ionomycin is added to equilibrate internal and external calcium in order to obtain the maximal Fluo-4 response. The data collected during these experiments were analyzed using Image J, Excel (Microsoft, WA) and Matlab (Mathworks, MA). Statistical significance was evaluated with Wilcoxon’s rank sum test.

## Results

### Structural homology and sequence alignment predict a PI(4,5)P_2_ binding site in TRPV1

Although the recently solved structure of a PI(4,5)P_2_ molecule located in an overlapping position with the capsaicin binding site of TRPV1 signifies an inhibitory role for the lipid, the existence of several TRP channel structures with alternate PI(4,5)P_2_ binding sites inspired us to use a homology based approach to scour the TRPV1 interaction surface for alternative binding sites [22, 26, 27]. Our previous MD study of the pH dependent interaction between PI(4,5)P_2_ and TRPV5 directed our search to a site proximal to the TRPV1 “inhibitory” phosphoinositide (Shown in **Figure 1A, B**, with dic8-PI(4,5)P_2_ bound in both TRPV1 and TRPV5 structures). In our simulation of TRPV5 and dic8-PI(4,5)P_2_, we examined the dynamic behavior of the lipid as it entered the binding site and determined that the inositol head group undergoes large movements until a distance of ~2 Å from the fully docked position. Notably, the headgroup rotates to accommodate an exchange of the 5’ phosphate interaction from R492 to R305 as the lipid shifts into a final position. Remarkably, both R492 and R305 in TRPV5 have an excess of histidine variants in the human population, which may be relevant to the pH dependent character of interacting with the 5’ phosphate (**Table S1**). Several aspects of the TRPV5 PI(4,5)P_2_ site became salient as we considered the same location in TRPV1: 1. The PI(4,5)P_2_ interacting residues of TRPV5 are identical or charge conserved in TRPV1 (**Figure 1A,B** highlighted); 2. The interacting TRPV1 residues are conserved across species, an exceptional change being found in horse with glutamine in place of H410 (**Figure S1**; see Discussion); 3. An “open pore” structure of TRPV1 bound with DkTx and RTX (PDB: 7L2M) brings all the aforementioned residues into closer alignment with the TRPV5 residues, thereby adopting the PI(4,5)P_2_ bound configuration of TRPV5 (**Figure 1C;** TRPV1 in “open” state is magenta). In **Figure 1D** we list the top four PI(4,5)P_2_ interacting residues that we identified in our TRPV5 binding MD study with relevant TRPV1 counterparts and their assigned topological domain in the channel. Of the listed residues, H410 of TRPV1 strays the furthest, chemically speaking, from its related arginine in TRPV5. Based on variant analysis, H410 is also an attractive residue for our studies since TRPV5-R305 has 21 histidine alternative alleles in the human population (**Table S1**) [28]. We were intrigued by the idea that TRPV1 evolved to have a pH sensitive side-chain since our MD study of the cognate residue R305 in TRPV5 had extensive interactions with the 5’ phosphate of PI(4,5)P_2_. Furthermore, we obtained binding energy profiles of PI(4,5)P_2_ with TRPV5 and discovered that, under nominal pH conditions, the 5’ phosphate of the inositol headgroup was more likely to be deprotonated than the 4’ phosphate as it interacted with TRPV5. This led us to speculate that PI(4,5)P_2_ interactions with TRPV5 were pH dependent and mediated through the electrostatic interactions between R305 and the 5’ phosphate of the lipid. We therefore hypothesized that the pH dependence of an interaction between PI(4,5)P_2_ and TRPV5 would transfer to TRPV1 if both channels shared a common PI(4,5)P_2_ binding site. To begin, we determined whether charge manipulation at the H410 site through mutagenesis gave predictable outcomes in TRPV1 function.

### Substitution with positively charged arginine at H410 increases TRPV1 activity

Structural data and our MD simulations of TRPV5 interactions with the 5’ phosphate of PI(4,5)P_2_ guided our decision to mutate H410 of TRPV1 to arginine to test if this enhances channel function. We observed a left shift in the dose-response curve of the TRPV1-H410R mutant relative to wild type as would be expected if the interaction between PI(4,5)P_2_ and channel led to a gain of function (**Figure 2B,E**). In contrast to our substitution of a positive charge at H410, our substitution of the neutral glutamine in the same position did not shift the dose-response curve significantly from wild-type (**Figure 2C,E**). Finally, we substituted an aspartic acid for H410 to imbue the site with a distinctly negative charge. Since the bath solutions used in our excised patch experiments are buffered at pH 7.2, we anticipated that the 5’ phosphate of PI(4,5)P_2_ could assume a deprotonated state with a negative charge [21]. Contrary to our prediction that swapping the charge in H410 would disrupt any interaction with PI(4,5)P_2_ and restrict channel function, we observed an increase in the sensitivity to capsaicin with H410D (**Figure 2D,E**). Furthermore, some of the H410D dose response data sets, which we did not use in our final analysis of capsaicin sensitivity, experienced an unusual regime of current run-up after 1 μM capsaicin was applied to the patch (**Figure S2**).The results obtained with H410D led us to speculate that this mutation may strongly interfere with PI(4,5)P2 interactions. Therefore, we ventured forward in two directions to ask: 1. Does the substitution to arginine at H410 in TRPV1 maintain the pH dependence of activity observed in TRPV5? 2. Do TRPV1-H410D channels retain functional properties that depend on PI(4,5)P_2_?

### R410 mutant maintains pH dependence of wild type TRPV1 but loss of charge in Q410 mutant abolishes pH dependence

In order to examine how the side chain of H410 would interact with a phosphate group of PI(4,5)P_2_, we compared the capsaicin responses of channel activity at pH 7.2 and 8.0. Without the histidine, pH dependence of the electrostatic interaction between the side chain of residue 410 and inositol phosphate would fall entirely onto the protonation state of the phosphate. We therefore predicted that if the H410 is mutated to arginine, deprotonation of PI(4,5)P_2_ under basic conditions would strengthen the interaction with the arginine, and a leftward shift of the dose-response curve suggests a positive effect of the lipid on capsaicin activity. At pH 8.0 the dose-response curve of wild type and H410R TRPV1 undergoes a leftward shit relative to capsaicin responses at pH 7.2 (**Figure 3**). To disrupt the pH dependence of an interaction between PI(4,5)P2 and TRPV1, we tested the dose-response behavior of TRPV1-H410Q at pH 8.0. We obtained dose response curves at pH 7.2 and 8.0 for TRPV1-H410Q that did not differ significantly and surmised that the reduction of interactions between H410Q and PI(4,5)P_2_ would account for no observed changes in channel activity (**Figure 3**). Our study of pH dependence of TRPV1 and mutations at the 410 position yields a pattern of results consistent with pH dependence observed in TRPV5 activity.

In summary, our mutagenesis and pH dependent study of position 410 support a model of interaction between PI(4,5)P_2_ and TRPV1 that enhances channel activity. This contradicts structural evidence from TRPV1 in closed state structures with PI(4,5)P_2_ molecules in the vanilloid site that are close enough to interact with H410. Our electrophysiology results from H410R and H410Q mutants an pH dependence of channel currents support the existence of an alternative, second site, which is approximated by the position of PI(4,5)P_2_ in a TRPV5 structure and which promotes channel opening. The one outlying piece of evidence in this scheme is the H410D mutant since it enhanced channel activity, but we postulated that the mutation may be responsible for disrupting an interaction with a PI(4,5)P_2_ molecule in the primary phosphoinositide site. Because H410 is implicated in both a role for inhibition through the primary, vanilloid site and potentiation at a secondary site, we propose that the side chain of H410 is free to interact with the PI(4,5)P_2_ in two distinct binding sites, which confers a central role for lipid recognition on this residue. We decided it would be best to proceed with experiments that establish the PI(4,5)P2 sensitivity in H410D because we hypothesized that the interaction is lost and the lipid is no longer associated with the channel.

### Current sensitivity to poly-lysine treatment remains intact in the TRPV1-H410D mutant and Mg^2+^ applied to the inner leaflet has a reduced effect on TRPV1-H410Q relative to wild type channel

Our unexpected TRPV1-H410D results led us to examine whether this channel’s sensitivity to phosphoinositides could be tested with methods that manipulate free phosphoinositide levels in the membrane. The effect of phosphoinositide signaling on plasma membrane receptors and channels is commonly evaluated by application of cationic solutions such as short-chain poly-lysines and aminoglycoside at the inner leaflet of membrane [29, 30]. We tested the effect of poly-lysine solutions (3μg/mL) on the inner leaflet of excised patches that express wild type or TRPV1-H410D. Consistent with previous poly-lysine experiments done with wild type TRPV1, the capsaicin activated currents are suppressed in combination with the poly-lysine[9]. We observed a similar current reduction in the H410D mutant and surmised that these channels remain sensitive to the effects of poly-lysine. Upon washout of the poly-lysine, but with capsaicin still in the solution, both mutant and wild type have different degrees of current recovery (**Figure 4A**). However, this remains compatible with known variability in poly-lysine washout[29]. In lieu of polylysine solutions, we used Mg^2+^ solutions that would sequester polyanionic lipids when applied but would still be expected to wash out in buffer[31]. Since TRPV1 current is selective for divalent cations over monovalent, we anticipated some current block with application of 50 mM Mg^2+^ solutions, which could be dependent on channel activity (**Figure 4B**). Nevertheless, our H410 mutants responded to capsaicin with similar or left-shifted dose-response profiles relative to wild type(**Figure 2**). We therefore predicted that current block at 1μM capsaicin would be similar across our tested constructs and additional changes in the current would result from the PI(4,5)P_2_ sequestering properties inherent to 50 mM Mg^2+^. Current inhibition was similar in both wild type and H410D. However, significantly less inhibition of current by Mg^2+^ was measured in H410Q than in wild type (**Figure 4**). If Mg^2+^ were effective at sequestering PI(4,5)P_2_ from channels that required the lipid for activation, we would predict that the currents in wild type and H410Q channels would be more impacted by the Mg^2+^ solution than H410D, which would not interact with PI(4,5)P_2_. Our result did not lend itself to a simple interpretation that depends on sequestration of PI(4,5)P_2_. However, we were intrigued by the alternative possibility that Po differs significantly for these channels under the capsaicin activation conditions of these experiments. Although the dose response profiles of H410D and H410Q indicated ~75% of full activation with 1 μM capsaicin (**Figure 2**), these macroscopic currents do not provide information about the absolute open probability of channels. Two indicators that supported this prospect were the unusual run-up in current that we observed in some experiments with H410D, which was more apparent at concentrations >0.3 μM capsaicin, and the notable current variance in H410Q (**Figure 4A (bottom), 4B(bottom) and Figure S2**). We decided to examine this phenomenon by measuring capsaicin dependent current run-up across several of our channel mutants.

**Figure 4.**
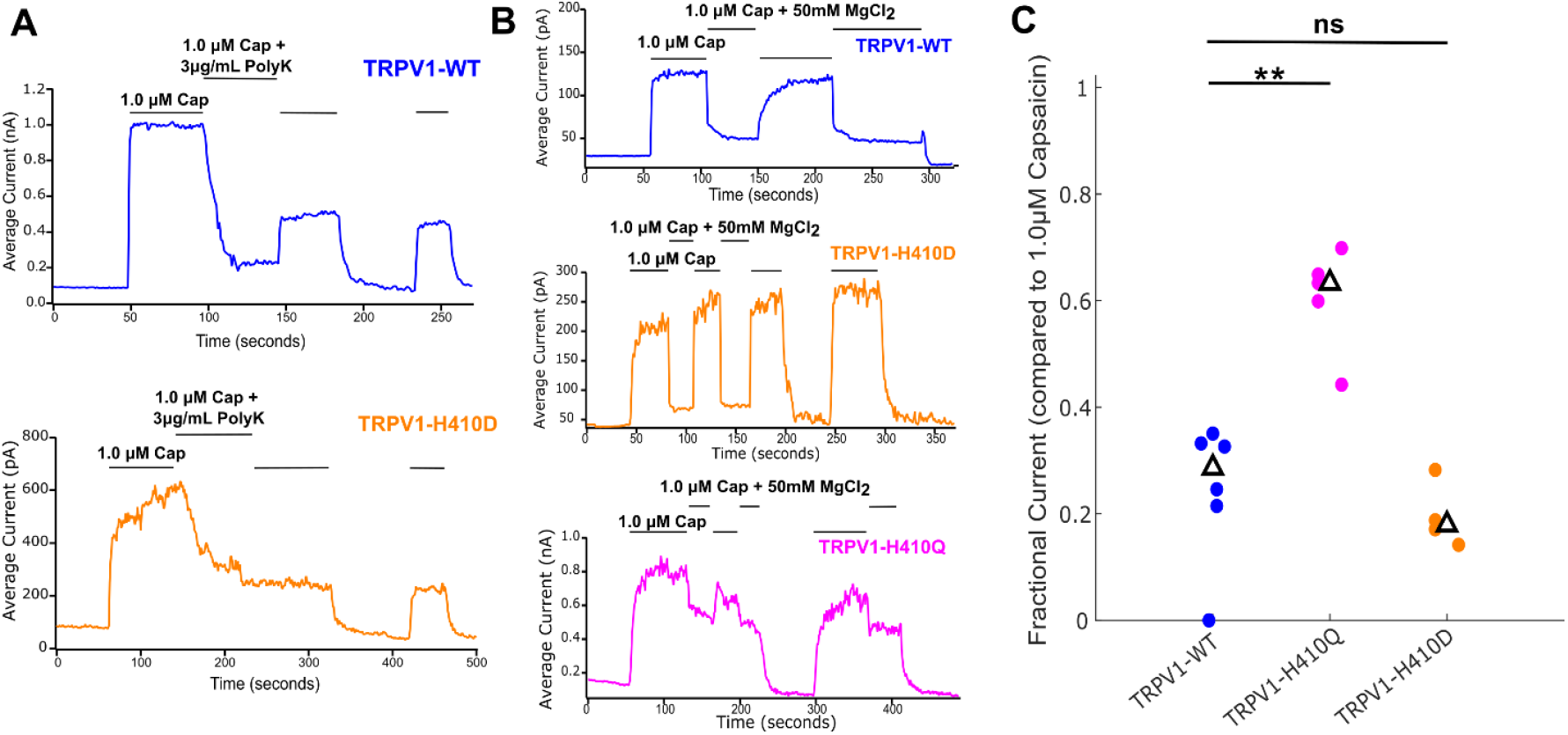
TRPV1 current sensitivity to Mg^2+^ applied to inner leaflet of excised membrane patches is altered in H410Q mutant. (A) Poly-lysine applied to TRPV1 wild type (top) and TRPV1-H410D (bottom) reduces capsaicin activated currents. (B) Effect of Mg^2+^ (50 mM) observed in representative, capsaicin activated currents from excised patches expressing TRPV1 wild type (top), TRPV1-H410D (middle), TRPV1-H410Q (bottom) channels. (C) Summary plot of fractional current with Mg^2+^ added to 1.0 μM capsaicin activation in wild type (WT), H410Q, and H410D TRPV1 channels. (‘**’: statistically significant at p = 0.0007; ‘ns’: not significant. Student’s two sample, two-sided t test)

### Prolonged capsaicin activation of the TRPV1 H410D mutant facilitates channel activity

We tested the concept that TRPV1 H410 mutant channels experience a time-dependent current run-up when activated by capsaicin. Our experiment consisted of two test exposures to 0.1 μM capsaicin before and after a prolonged treatment (~2 minutes) with 3.0 μM capsaicin. A ratio between the amplitude of current in the second 0.1 μM test pulse compared to the first served as a metric of current run-up induced by the 3.0 μM capsaicin. We selected the specific concentration of capsaicin during the prolonged treatment based on its placement in the 75%-100% range of full activation observed in wild type, H410D, and H410Q channels, and it exceeding the concentration needed for prior occurrences of run-up with H410D (**Figure 2, Figure 4A, bottom** and **Figure S2)**. When we tested the facilitation effect in TRPV1 wild type, we obtained a distribution of responses to 0.1 μM capsaicin after 2 minutes of activation with 3.0 μM capsaicin that did not appear to change from initial activity with 0.1 μM capsaicin (**Figure 5A,E**). We included the double mutant TRPV1 H410R/K688R with mutations to cognate position in the TRPV5-conserved PI(4,5)P_2_ binding site to examine differences in current run-up that would possibly arise from enhancing an interaction with PI(4,5)P_2_ relative to wild type **(Figure 1C, Figure S3**), but we did not observe a significant change after prolonged exposure to 3.0 μM capsaicin with this construct (**Figure 5B,E**). We observed a significant change to higher currents with the 2^nd^ capsaicin pulse in H410D (**Figure 5C,E**), and although a trend to higher currents appeared in the 2^nd^ pulse with H410Q, it was not significant at the p < 0.05 level (**Figure 5D,E**). These results suggest that a large pool of H410D channels are unresponsive to low capsaicin concentrations, possibly remaining in a quiescent state. Higher concentrations of capsaicin (>0.1 μM) are then effective at bringing these “silent” channels into activated states. Two properties in H410D current run-up experiments lead us to believe that channel facilitation is responsible for the effect: (1) the 2^nd^ pulse current in H410D approaches a stable end-point that is greater than the current of the 1^st^ pulse (2) when facilitation occurred in dose response experiments, a return to capsaicin activated currents after washout proceeded with rapid kinetics (**Figure S2**). It was of some relevance that the study by Arnold and colleagues used excess capsaicin to displace PI molecules in reconstituted TRPV1 vesicles which were used to furnish their study with an open state channel that has neither capsaicin nor PI bound in the primary binding site [17]. This suggested that an active state of the channel, facilitated by loss of PI, is maintained after removal of the capsaicin and that the channel pore does not easily return to a closed state without a phosphoinositide in the primary site. H410D would likely interfere with PI(4,5)P_2_ interactions in the vanilloid site but not necessarily a PI. Our facilitation results can, therefore, be explained by a model in which capsaicin displaces the PI molecule residing in the vanilloid site of a resting TRPV1-H410D channel, and the channel remains in a facilitated, active state long after capsaicin is removed (**Figure 2D, 5C, long recovery to closed state**). Based on our results and the structural evidence that PI is often identified in the primary phosphoinosiide site, we hypothesized that the H410D mutation selects for PI over PI(4,5)P_2_ in the primary, vanilloid site, casting the side-chain of H410 in a role for phosphoinositide recognition in the primary site.

**Figure 5.**
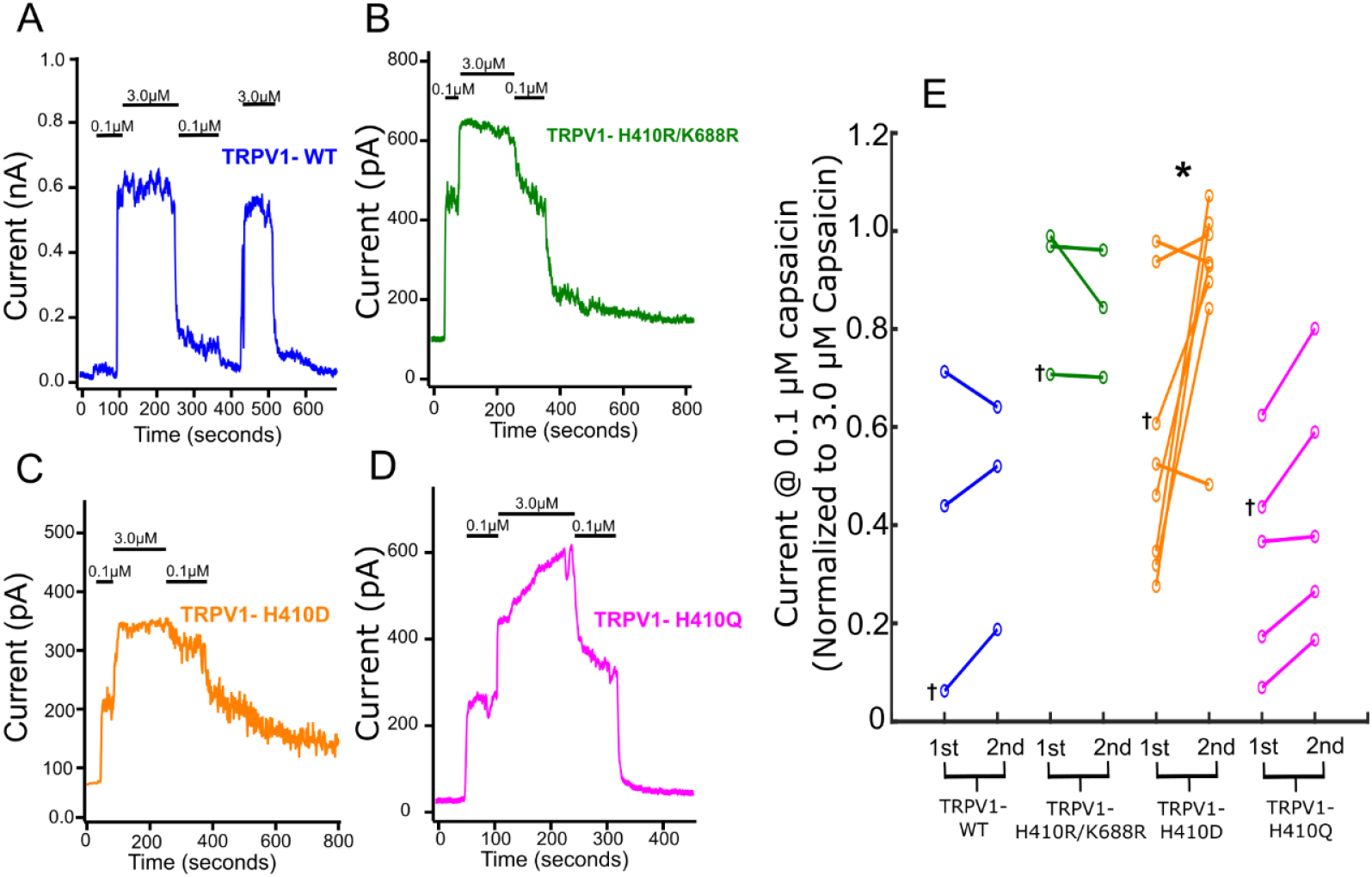
High concentrations of capsaicin promote current run-up in excised patches expressing TRPV1-H410D. (A-D) TRPV1 wild type and TRPV1-H410 mutant representative traces of current response to a baseline of 0.1 µM capsaicin followed by perfusion of 3.0 µM capsaicin for 120-180 seconds, then subsequent test pulse with 0.1 µM capsaicin. (E) Summary data of the 0.1 µM capsaicin responses 1^st^, before the 3.0 µM perfusion, and 2^nd^, after the 3.0 µM perfusion shown on a paired dot plot. The 1^st^ and 2^nd^ values were normalized by dividing the final current in the 0.1 µM section by the maximal current of the 3.0 µM capsaicin response. The daggers represent the summary data points that correspond to the representative traces in A-D. A one sample Student’s t test centered at 1, was performed to compare the ratios of the 2^nd^ 0.1 µM currents to the 1^st^ 0.1 µM capsaicin currents and observed the following p-values TRPV1 wild type (p=0.4021) TRPV1-H410R/K688R (p= 0.3598), TRPV1-H410D (p= 0.0345), and TRPV1-H410Q (p= 0.0920). (*; statistically significant at p=0.0345. Student’s one sample centered at 1, two-tailed t test)

### Loss of the PI(4,5)P_2_ associated regulation in TRPV1 H410D channels

Our interest in knowing whether the H410D mutant still interacts with PI(4,5)P_2_ led us to examine next whether the H410D mutant is still regulated by PI(4,5)P_2_. The controversial role of PI(4,5)P_2_ regulation in TRPV1 is well-described by two types of experiments, each tested at opposite ends of the concentration range of capsaicin activation. In the first type of experiment that tests for PI(4,5)P_2_ inhibition, the resident PI(4,5)P_2_ in the plasma membrane (reported to be in the range of 1-2% mol fraction) is depleted by means of an antibody or reconstituted phospholipase C (PLC) and non-saturating capsaicin currents are shown to increase [8]. The second type of experiment tests for potentiation by reducing TRPV1 currents attributable to near-saturating concentrations of capsaicin with the application of PI(4,5)P_2_-specific binding proteins or phosphatase activity against the 5’ phosphate of the inositol ring [9, 12]. We attempted to address whether these two aspects of channel regulation are intact in TRPV1-H410D. If PI(4,5)P_2_ cannot interact with the TRPV1 H410D mutant, we predicted that both (1) inhibition by PI(4,5)P_2_ at low activation thresholds and (2) potentiation by PI(4,5)P_2_ of sustained currents under near-saturating capsaicin conditions would be lost. To test whether the PI(4,5)P_2_ inhibition modality of TRPV1H410D is intact, we opted to manipulate PI(4,5)P_2_ levels with a rapamycin inducible phosphatase recruitment tool in intact cells and follow TRPV1 activity with calcium imaging[12]. The rapamycin inducible PI(4,5)P_2_/PIP phosphatase recruitment system incorporates the Sac1 and inositol 5’-phosphate phosphatases in a membrane recruitable fusion protein called Pseudojanin (PJ), which rapidly converts PI(4,5)P_2_ and PI(4)P into PI. The PJ double phosphatase is fused to an FKB protein, which interacts with the FRB domain when bound by rapamycin. A separate construct expresses the Lyn_11_-FRB fusion protein which is targeted to the plasma membrane to ensure efficient translocation of the PJ to the plasma membrane when rapamycin is perfused over the intact cells. In our experiment we perfused 1 μM rapamycin over cells that expressed the PJ-rapamycin phosphatase system in combination with the TRPV1 channel to test whether loss of PI(4,5)P2 would remove phosphoiniositide based inhibition over TRPV1 and lead to spontaneous channel activation. Even without the activating presence of capsaicin, rapamycin induction of PJ activity and conversion of PI(4,5)P_2_ and PI(4)P2 into PI elicited a rapid and transient calcium response in TRPV1 wild type expressing cells (**Figure 6A**). If the TRPV1 bound PI(4,5)P_2_ is coupled to equilibria of membrane PI(4,5)P_2_, we speculate that loss of inhibition was achieved by simply reducing cellular pools of PI(4,5)P_2_ and the effect of mass action on PI(4,5)P2 binding to the primary, vanilloid site of the channel. In sharp contrast to wild type, the H410D mutant showed no responses to rapamycin induction although sensitivity to capsaicin remained intact across cells that expressed the PJ system (**Figure 6B, compare H to N**). Evidently, rapid membrane removal of PI(4,5)P_2_ only affected wild type channels, implicating H410D as a mutation that abrogates TRPV1 inhibition by P(4,5)P2. This result corroborates previous studies that remove inhibition of TRPV1 by depleting the plasma membrane of PI(4,5)P_2_ either by sequestering with a PI(4,5)P_2_ antibody or enzymatically degrading PI(4,5)P_2_ with PI-PLC. Because the second type of PI(4,5)P_2_-based regulation over TRPV1 is observed with high activity at near-saturating capsaicin concentrations, we tested TRPV1 wild type and H410D responses with whole cell electrophysiology. Currents from cells expressing the PJ-rapamycin recruitment system and TRPV1 constructs were first elicited by high capsaicin concentrations (3 μM) before switching into a solution that contained 1 μM rapamycin and 3 μM capsaicin (**Figure 6P,Q**). Cells expressing wild type channel responded to the rapamycin/capsaicin solution with a reduction in overall current as potentiation by PI(4,5)P_2_ is lost while cells expressing the H410D TRPV1 construct had currents that remained high and unaffected **(Figure 6P,Q,R)**. The whole cell current response with wild type TRPV1 corroborates previous experiments performed with membrane recruited phosphatases specific for PI(4,5)P_2_. The novel result with H410D confirms that current potentiation by PI(4,5)P_2_ is not possible in this mutant. The simplest explanation for a lack of PI(4,5)P_2_ sensitivity in H410D under both experimental paradigms is that the mutant does not interact with PI(4,5)P_2_ at either the primary inhibitory site or the TRPV5-conserved potentiating site. Moreover, the whole cell current experiment vindicates PI(4,5)P_2_ as a bonified potentiator of current in wild type channels. Although PI was generated in excess as a product of PJ-rapamycin activity, conversion of PI(4,5)P_2_ to PI did not interfere with the whole cell currents measured in the H410D mutant which would still be susceptible to PI’s competitive displacement of capsaicin at the vanilloid site.

**Figure 6.**
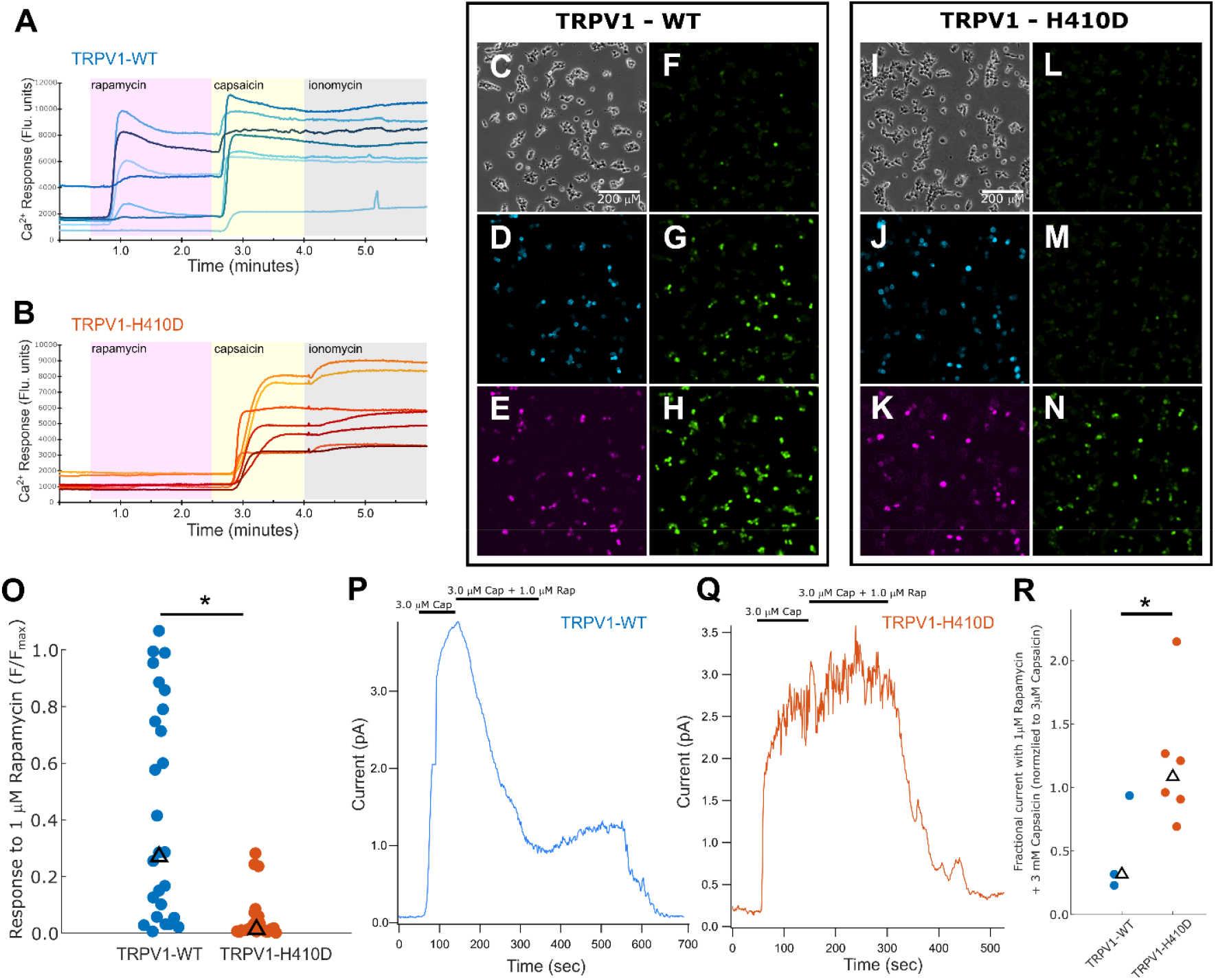
Loss of PI(4,5)P_2_ by activation of PJ-rapamycin system both removes inhibition and reduces potentiation in wild type (WT) TRPV1 but not TRPV1-H410D. (A) Fluorescence responses from Fluo-4 measuring Ca^2+^ influx in intact cells co-transfected with wild type TRPV1, mRFP-FKBP-Pseudojanin (PJ), and Lyn11-FRB-CFP. Rapamycin (1 μM) is added after 30s and perfusion is changed over to 0.1 μM capsaicin at the 2 minute, 30 second mark. Ionomycin (5 μM) added at 4 minutes to establish maximal fluorescence response (Fmax). (B) Identical experiment as described in panel (A) but with TRPV1-H410D instead of wild type channel in the co-transfection. (C) bright field image of cells analyzed in panel (A). (D) CFP epi-fluorescence image of cells analyzed in panel (A) to reveal expression of Lyn_11_-FRB-CFP construct. (E) RFP epi-fluorescence image of cell analyzed in panel (A) to reveal expression of PJ. (F-H) Panels show Ca^2+^ signal response with GFP epi-fluorescence image of cells analyzed in panel (A) at 0, 1 minute, 3 minutes, which show responses with: no stimulation, activation with rapamycin (1 μM), activation with capsaicin (0.1 μM). (I) bright field image of cells analyzed in panel (B). (J) CFP epi-fluorescence image of cells analyzed in panel (B) to reveal expression of Lyn_11_-FRB-CFP construct. (K) RFP epi-fluorescence image of cell analyzed in panel (B) to reveal expression of PJ. (L-N) Panels show Ca2+ signal response with GFP epi-fluorescence image of cells analyzed in panel (B) at 0, 1 minute, 3 minutes, which show responses with: no stimulation, activation with rapamycin (1 μM), activation with capsaicin (0.1 μM). (O) Summary plot of TRPV1 wild type and TRPV1-H410D fluorescence responses to rapamycin (1 μM) normalized to max fluorescence measured with ionomycin (Fmax) (‘*’: statistically significant at p=0.00002 with Wilcoxon rank sum test). (P) Representative trace of TRPV1 wild type (WT) currents in whole-cells expressing PJ-rapamycin system and activated with 3 μM capsaicin followed by switch to 1 μM rapamycin + 3 μM capsaicin. Switching the amplifier to lower gain sensitivity at ~100 seconds introduced the short flat segment in currents. (Q) Representative trace of TRPV1-H410Q currents in whole-cells expressing PJ-rapamycin system and activated with 3 μM capsaicin followed by switch to 1 μM rapamycin + 3 μM capsaicin. (R) Summary plot of fraction of current observed after application of 1 μM rapamycin + 3 μM capsaicin compared to 3 μM capsaicin alone in cells expressing TRPV1 wild type or TRPV1-H410D alongiside PJ-rapamycin system. (*; statistically significant at p=0.038. Student’s two sample, one-sided t test)

## Discussion

Regulation of TRPV1 function encapsulates a complex network of cell signaling pathways involving calcium, phosphorylation, and lipids, to name a few modalities. For the peripheral neurons that endogenously express this channel, tight control over TRPV1’s regulatory apparatus is necessary to accurately transmit signals from initial stimuli to downstream synapses. Since the discovery of TRPV1 by expression cloning over 25 years ago, researchers have made important contributions to our understanding of this channel’s activation properties and regulatory mechanisms. Lipid regulation by phosphoinositides remains one of the more controversial topics in the history of TRPV1 studies. Beginning in the early 2000s, studies were published that appeared to form a back-and-forth debate on whether PI(4,5)P_2_ would act as an inhibitor or cofactor of TRPV1. Now, with two decades of functional studies and a large assortment of biological structures to assist our process of understanding TRPV1, we have made critical progress towards clarifying lipid regulation over this channel. Certainly, PI is frequently bound to closed channel states in well-resolved structures, and the site where it binds partially overlaps with the vanilloid binding site, situating the PI as a competitive inhibitor. However, structures were only recently solved with PI(4,5)P_2_ located in the same position as PI, and the elaborate experimental process of first removing the endogenous PI with capsaicin before introducing recombinant PI(4,5)P_2_ to obtain these structures inspired us to consider what the mechanistic underpinnings of this exchange were[17]. Moreover, several other groups proposed that more than one phosphoinositide binding site existed in TRPV1[4, 20, 32]. Finally, what has remained at the focus of our interests in these matters are the functional studies that identified PI(4,5)P_2_ as an inhibitor that still need to be reconciled with data that shows how the molecule potentiates currents.

Our model of TRPV1 regulation by lipids is defined by two sites that interact across the residue H410 (**Figure 7**). Functional studies with the H410 position showed that an enhancement in function could be achieved by strengthening the electrostatic interaction between PI(4,5)P_2_ and channel with the H410R mutation (**Figures 2**,**3**). A model that only supports binding of phosphoinositides at the primary, inhibitory site is not compatible with this result, but a second phosphoinositide binding site that maintains lipid interactions with H410 would complement our findings. Importantly, preserving the H410 interaction with the phosphoinositide in either site necessitates a mutually exclusive phosphoinositide binding paradigm (**Figures 5**,**6**). We speculate that PI(4,5)P_2_ cannot occupy both sites simultaneously, and the H410 residue would shuttle between sites to accompany the translocation of the phosphoinositide (**Figure 7, center**). The compact and unique molecular architecture of capsaicin allows it to bind in the primary phosphoinositide site while PI(4,5)P_2_ interacts with the secondary site to stabilize the open-state channel. This binding arrangement could serve as a general model for how smaller, lipophilic molecules serve as partial agonists in TRPV1 activation[5].

**Figure 7.**
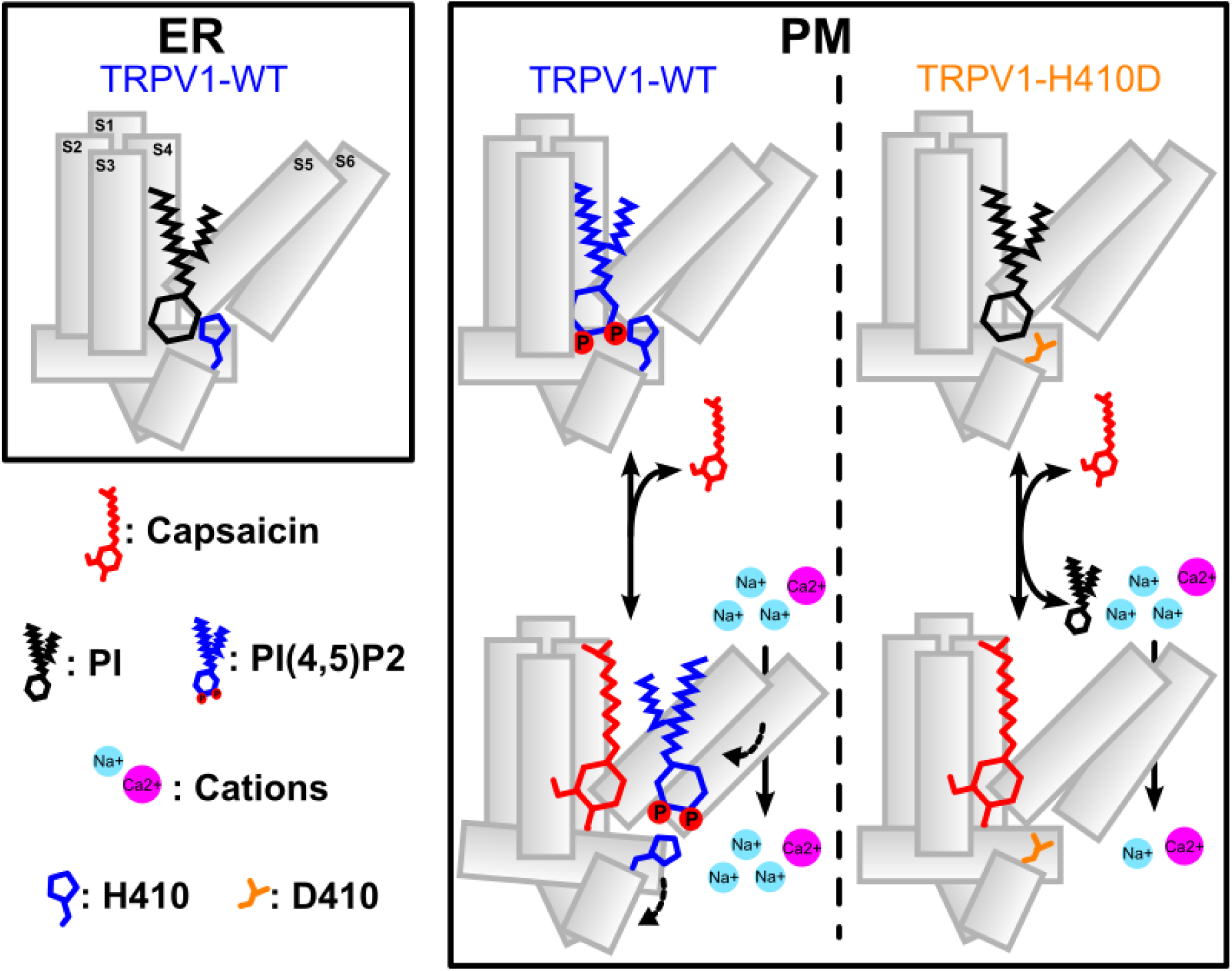
Proposed model for two PI(4,5)P_2_ binding sites in TRPV1. (Left) TRPV1 - shown as a single subunit - expressed in the ER is bound by PI in the primary, inhibitory binding site. (Center) TRPV1 wild type channel exchanges PI for PI(4,5)P_2_ in the plasma membrane (PM). Capsaicin displaces PI(4,5)P_2_ from the primary binding site and H410 coordinates the transfer of PI(4,5)P_2_ into the second, “front porch” site, which favors an open state channel. The mutually exclusive binding of phosphoinositides at the primary, vanilloid and secondary, front porch, site is sustained in the lower panel because capsaicin is a smaller lipophilic molecule than a phosphoinositide.(Right) TRPV1-H410D retains the PI from the ER when residing in the PM. Capsaicin displaces the PI from the primary binding site when channel gates, but the D410 residue cannot accommodate the PI(4,5)P_2_ in the vanilloid site nor the “front porch” site, which precludes TRPV1-H410D from sensitivity to manipulations of PI(4,5)P_2_ levels.

It is well-established that different intracellular membranes have characteristic phosphoinositide compositions that serve as molecular markers[33, 34]. PI, is more broadly distributed across the cell since it forms the basis of generating all the singly phosphorylated phosphoinositides, giving different intracellular membranes a means to build molecular identity with membrane bound kinases and phosphatases[35]. Nevertheless, recent studies have deduced that the amount of PI in the plasma membrane is considerably lower than in the golgi or ER [36, 37]. PI(4,5)P_2_, on the other hand, is found at a much higher mol fraction in the plasma membrane (~1-2%) compared with other cellular compartments, which implies a discernable shift in the ratio of PI to PI(4,5)P_2_ experienced by a membrane associated protein as it is trafficked from the site of expression in the ER to the plasma membrane. [38, 39]. We speculate that our H410D mutant exhibited a complete lack of PI(4,5)P_2_ dependent phenomena because the channel remains associated with PI instead of exchanging it for PI(4,5)P2 found in the plasma membrane. PI bound TRPV1 is, therefore, likely to represent the state of TRPV1 found in intracellular membranes such as the ER or a channel that is newly inserted into the plasma membrane.

A convenient description of TRPV1 maturation and how the PI is exchanged for PI(4,5)P_2_ in the plasma membrane follows from our model (**Figure 7**). Once in the plasma membrane, TRPV1 experiences a conformational change, perhaps even given some impetus by thermal activation of the channel, that releases PI from the primary site and moves H410 into a position that is prone to interact with PI(4,5)P_2_, which is the TRPV5-conserved site that we have termed the “front porch”. Once the channel “catches” PI(4,5)P_2_ in the front porch, the side chain of H410 would maintain an interaction with the 5’ phosphate of the inositol head group and accompany the lipid back into the primary phosphoinositide site that promotes channel closure. Ultimately, movement of a lipid out of the primary, vanilloid site removes channel inhibition[17], ensuring that the front porch binding site is activating with a higher affinity for PI(4,5)P_2_ than PI thus guaranteeing a rapid exchange process for the phosphosinositides (**Figure 7, center**). In fact, our initial, functional characterization of the H410R mutant served as a prescient indicator for our model since the only capsaicin concentration with a significantly increased channel activity over wild type was at 1.0 μM, which would coincide with channels populating a state that binds PI(4,5)P_2_ at the front porch site (**Figure 2E, teal**). At lower concentrations of capsaicin, the majority of channel bound PI(4,5)P_2_ is still located at the primary site, where the lipid cannot increase channel activity by interacting favorably with the substituted arginine. Moreover, structural data (**Figure 1C**) and the mutants H410R and H410Q with their relevant pH dependent experiments (**Figure 2,3**) assign an activating and not inhibitory status to the front porch site. Finally, the prediction of the front porch site was made possible by structural comparison to TRPV5, which shares only 30.6% sequence homology with TRPV1. This then beckons the question of whether our description of phosphoinositide regulation in TRPV1 could be a general mechanism for all channels in the TRPV subfamily. In TRPV3, for example, currents activated by 2-APB or camphor appear to be facilitated by prolonged exposure to agonist in the same way as TRPV1-H410D[40, 41]. It is therefore plausible that the facilitation effect in both channels has a common mechanistic origin through a lipid occupying the vanilloid site, which can be competed away with a small lipophilic agonist

One important conclusion we can draw from our experiments is that the mature form of closed state TRPV1 in plasma membrane is complemented with PI(4,5)P_2_ in the primary, vanilloid site. Because of this, structural studies of TRPV1 that incorporate lipids need to interpret the presence of lipid and assignment of channel state with care. TRPV1 with only PI in the primary phosphoinositide site would not represent a typical closed state channel found in the plasma membrane, where its activity is of biological concern. Moreover, our functional experiments with TRPV1-H410D set an important standard for TRPV1 structural studies and may necessitate reconstitution efforts with PI(4,5)P_2_ for structural models to provide an accurate description of channel gating.

Although the charge neutralization mutation for H410 was possible with glutamine or asparagine, our choice of H410Q was not arbitrary. Initial sequence alignments brought to light that equine TRPV1 has a glutamine at this position, and our results may offer some insight into the relevance of this substitution. Horses are notable for their incredible athleticism and metabolic activity [42]. Accumulation of metabolites in serum such as lactate can exceed twice the amount in humans, but probably more profound (and relevant to discussions about TRPV1) is the temperature increase in horse muscle, which can exceed 42 C in the exercising animal[43]. It is therefore not altogether surprising that the H410Q mutation may be a unique adaptation to the biological character of the horse that mitigates its extreme physiological experiences. Firstly, the H410Q channel appears to have a reduced sensitivity to pH changes which may be an adaptation to the rapid lactic acid production in exercise, and secondly, the channels exhibit some of the slow facilitation of current that may be important to keep TRPV1 activity muted if muscle temperature routinely exceeds 40 C during exercise. An important future study would be to evaluate to what extent horse TRPV1 may undergo a facilitation process, for which we only observed a trend in H410Q of this study, and we would want to determine if the ratio of PI and PI(4,5)P_2_ in plasma membrane bound H410Q channels is fundamentally different from channels that are otherwise histidine at this position.

## Supplementary Data

### Supplemental Figures

**Figure S1. Sequence alignments of TRPV1 and TRPV5**. Red boxes outline PI(4,5)P_2_ interacting residues (numbering is rat TRPV1) identified in Figure 1D. Alignment sequence IDs: *H. sapiens* TRPV1, NM_080704.4; *R. norvegicus* TRPV1, NM_031982.1; *C. familiaris* TRPV1, NM_001003970.1; *C. porcellus* TRPV1: NM_001172652.1; *E. caballus* TRPV1, XM_014727972.3; *H. sapiens* TRPV5, NM_019841.7

**Figure S2. Examples of current instability in two dose response curves with expression of TRPV1-H410D in excised patches**. (A) Dose response experiment of TRPV1-H410D with stable currents < 0.3 μM capsaicin before run-up in currents through 5.0 μM capsaicin. Washout of capsaicin demonstrates that excised patch still has intact giga-seal. High activity currents elicited with 5.0 μM capsaicin exhibit notable variance in the current measurement. (B) Dose response experiment with current instability detected in 0.03 μM capsaicin and eventual run-up in currents with perfusion of capsaicin across concentration range. Washout of capsaicin is shown to demonstrate intact patch seal.

**Figure S3. TRPV1-H410R/K688R currents have a left shifted, capsaicin-activated dose-response profile compared to TRPV1 Wild type currents**. (A-B) Representative traces of capsaicin dose response experiments with TRPV1-H410R/K688R. Solution changes with different concentrations of capsaicin given in μM is depicted above the traces. (C) Summary of dose response data for TRVPV1-H410R/K688R show in in panels A and B compared with dose-response profile of TRPV1-WT. The TRPV1-WT capsaicin dose response profile was adopted from Figure 2E. Hill equation fit to the average of the TRVPV1-H410R/K688R capsaicin dose response experiments was used to acquire profile and EC_50_ values. Graph A corresponds to the triangles and graph B corresponds to the circle markers on this summary plot.

### Supplemental Tables

**Table S1. TRPV1 and TRPV5 genomic variants of interest**. Residues of interest to this study are listed with their associated allelic variants in the human population. Data are collected from 4.10 dataset in the Gnomad database (See methods)

## Acknowledgments

We thank Dr. Andrés Jara-Oseguera and Dr. Marcel Goldschen-Ohm for their helpful discussions and reading of the manuscript. Eric Senning wishes to thank his student researchers, whose hard work, perseverance and patience with him made this project possible. This work was supported by start-up funds from the University of Texas at Austin (E.N.S.) and the NSF-MCB grant no. 2129209 (E.N.S.).

## References

1. Caterina, M.J., et al., The capsaicin receptor: a heat-activated ion channel in the pain pathway. Nature, 1997. 389(6653): p. 816–24.

2. Davis, J.B., et al., Vanilloid receptor-1 is essential for inflammatory thermal hyperalgesia. Nature, 2000. 405(6783): p. 183–7.

3. Rosenbaum, T. and L.D. Islas, Molecular Physiology of TRPV Channels: Controversies and Future Challenges. Annu Rev Physiol, 2023. 85: p. 293–316.

4. Rohacs, T., Phosphoinositide Regulation of TRP Channels: A Functional Overview in the Structural Era. Annu Rev Physiol, 2023.

5. Hwang, S.W., et al., Direct activation of capsaicin receptors by products of lipoxygenases: endogenous capsaicin-like substances. Proc Natl Acad Sci U S A, 2000. 97(11): p. 6155–60.

6. Morales-Lázaro, S.L., et al., Structural determinants of the transient receptor potential 1 (TRPV1) channel activation by phospholipid analogs. J Biol Chem, 2014. 289(35): p. 24079–90.

7. Diver, M.M., et al., Sensory TRP Channels in Three Dimensions. Annu Rev Biochem, 2022. 91: p. 629–649.

8. Chuang, H.H., et al., Bradykinin and nerve growth factor release the capsaicin receptor from PtdIns(4,5)P2-mediated inhibition. Nature, 2001. 411(6840): p. 957–62.

9. Klein, R.M., et al., Determinants of molecular specificity in phosphoinositide regulation. Phosphatidylinositol (4,5)-bisphosphate (PI(4,5)P2) is the endogenous lipid regulating TRPV1. The Journal of biological chemistry, 2008. 283(38): p. 26208–16.

10. Senning, E.N., et al., Regulation of TRPV1 ion channel by phosphoinositide (4,5)-bisphosphate: the role of membrane asymmetry. J Biol Chem, 2014. 289(16): p. 10999–1006.

11. Ufret-Vincenty, C.A., et al., Mechanism for phosphoinositide selectivity and activation of TRPV1 ion channels. J Gen Physiol, 2015. 145(5): p. 431–42.

12. Hammond, G.R., et al., PI4P and PI(4,5)P2 are essential but independent lipid determinants of membrane identity. Science, 2012. 337(6095): p. 727–30.

13. Cao, E., et al., TRPV1 channels are intrinsically heat sensitive and negatively regulated by phosphoinositide lipids. Neuron, 2013. 77(4): p. 667–79.

14. Kwon, D.H., et al., Heat-dependent opening of TRPV1 in the presence of capsaicin. Nat Struct Mol Biol, 2021. 28(7): p. 554–563.

15. Sánchez-Hernández, R., et al., Structural basis of the inhibition of TRPV1 by analgesic sesquiterpenes. Proc Natl Acad Sci U S A, 2025. 122(29): p. e2506560122.

16. Zhang, K., D. Julius, and Y. Cheng, Structural snapshots of TRPV1 reveal mechanism of polymodal functionality. Cell, 2021. 184(20): p. 5138–5150.e12.

17. Arnold, W.R., et al., Structural basis of TRPV1 modulation by endogenous bioactive lipids. Nat Struct Mol Biol, 2024. 31(9): p. 1377–1385.

18. Kwon, D.H., et al., Vanilloid-dependent TRPV1 opening trajectory from cryoEM ensemble analysis. Nat Commun, 2022. 13(1): p. 2874.

19. Nadezhdin, K.D., et al., Extracellular cap domain is an essential component of the TRPV1 gating mechanism. Nat Commun, 2021. 12(1): p. 2154.

20. Yazici, A.T., et al., Dual regulation of TRPV1 channels by phosphatidylinositol via functionally distinct binding sites. J Biol Chem, 2021. 296: p. 100573.

21. Fathizadeh, A., E. Senning, and R. Elber, Impact of the Protonation State of Phosphatidylinositol 4,5-Bisphosphate (PIP2) on the Binding Kinetics and Thermodynamics to Transient Receptor Potential Vanilloid (TRPV5): A Milestoning Study. J Phys Chem B, 2021. 125(33): p. 9547–9556.

22. Hughes, T.E.T., et al., Structural insights on TRPV5 gating by endogenous modulators. Nat Commun, 2018. 9(1): p. 4198.

23. Gouy, M., et al., Seaview Version 5: A Multiplatform Software for Multiple Sequence Alignment, Molecular Phylogenetic Analyses, and Tree Reconciliation. Methods Mol Biol, 2021. 2231: p. 241–260.

24. Wulffraat, G.C., et al., Missense variant analysis in the TRPV1 ARD reveals the unexpected functional significance of a methionine. PLoS One, 2025. 20(9): p. e0331224.

25. Mott, T.M., et al., Fluorescence labeling strategies for cell surface expression of TRPV1. J Gen Physiol, 2024. 156(10).

26. Yin, Y., et al., Mechanisms of sensory adaptation and inhibition of the cold and menthol receptor TRPM8. Sci Adv, 2024. 10(31): p. eadp2211.

27. Cai, R., et al., Autoinhibition of TRPV6 Channel and Regulation by PIP2. iScience, 2020. 23(9): p. 101444.

28. Mott, T.M., et al., Mutagenesis studies of TRPV1 subunit interfaces informed by genomic variant analysis. Biophys J, 2023. 122(2): p. 322–332.

29. Hilgemann, D.W., Local PIP(2) signals: when, where, and how? Pflugers Arch, 2007. 455(1): p. 55–67.

30. Zhang, H., et al., PIP(2) activates KCNQ channels, and its hydrolysis underlies receptor-mediated inhibition of M currents. Neuron, 2003. 37(6): p. 963–75.

31. Suh, B.C. and B. Hille, Electrostatic interaction of internal Mg2+ with membrane PIP2 Seen with KCNQ K+ channels. J Gen Physiol, 2007. 130(3): p. 241–56.

32. Poblete, H., et al., Molecular determinants of phosphatidylinositol 4,5-bisphosphate (PI(4,5)P2) binding to transient receptor potential V1 (TRPV1) channels. J Biol Chem, 2015. 290(4): p. 2086–98.

33. Di Paolo, G. and P. De Camilli, Phosphoinositides in cell regulation and membrane dynamics. Nature, 2006. 443(7112): p. 651–7.

34. Posor, Y., W. Jang, and V. Haucke, Phosphoinositides as membrane organizers. Nat Rev Mol Cell Biol, 2022. 23(12): p. 797–816.

35. De Craene, J.O., et al., Phosphoinositides, Major Actors in Membrane Trafficking and Lipid Signaling Pathways. Int J Mol Sci, 2017. 18(3).

36. Zewe, J.P., et al., Probing the subcellular distribution of phosphatidylinositol reveals a surprising lack at the plasma membrane. J Cell Biol, 2020. 219(3).

37. Pemberton, J.G., et al., Defining the subcellular distribution and metabolic channeling of phosphatidylinositol. J Cell Biol, 2020. 219(3).

38. Balla, T., Phosphoinositides: tiny lipids with giant impact on cell regulation. Physiol Rev, 2013. 93(3): p. 1019–137.

39. Cullen, P.J. and J.G. Carlton, Phosphoinositides in the mammalian endo-lysosomal network. Subcell Biochem, 2012. 59: p. 65–110.

40. Peier, A.M., et al., A heat-sensitive TRP channel expressed in keratinocytes. Science, 2002. 296(5575): p. 2046–9.

41. Xu, H., et al., TRPV3 is a calcium-permeable temperature-sensitive cation channel. Nature, 2002. 418(6894): p. 181–6.

42. Rivero, J.L. and E.W. Hill, Skeletal muscle adaptations and muscle genomics of performance horses. Vet J, 2016. 209: p. 5–13.

43. Poole, D.C. and H.H. Erickson, Highly athletic terrestrial mammals: horses and dogs. Compr Physiol, 2011. 1(1): p. 1–37.

