## Supplemental Data for "A single residue determines regulation of TRPV1 by phosphoinositides"

Supplemental Figures

Figure S1

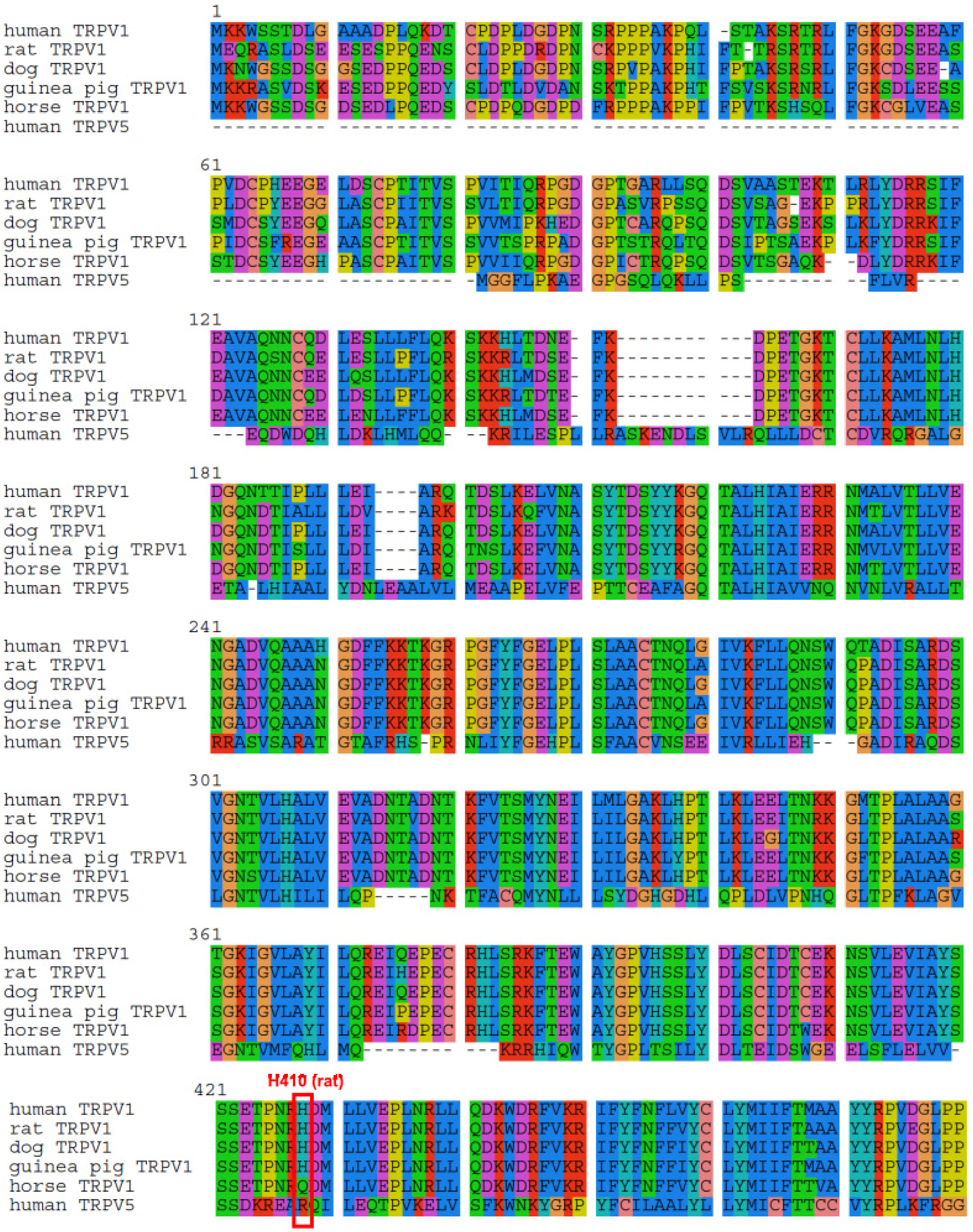

```
                     481
human TRPV1          FK-------- --------ME -KTGDYFRVT GEILSVLGGV YFFFRGIQYF LQRRPS---M
rat TRPV1            YK-------- --------LK NTVGDYFRVT GEILSVSGGV YFFFRGIQYF LQRRPS---L
dog TRPV1            YK-------- --------LK HTVGDYFRVT GEILSVLGGV YFFFRGIQYF LQRRPS---L
guinea pig TRPV1     YK-------- --------MK NTVGDYFRVT GEILSVIGGF HFFFRGIQYF LQRRPS---V
horse TRPV1          FK-------- --------LN -TVGDYFRVT GEILSVSGGV YFFFRGIQYF LQRRPS---L
human TRPV5          NRTHSRDITI LQQKLLQEAY ETREDIIRLV GELVSIVGAV IILLLEIPDI FRVGASRYFG

                     541
human TRPV1          KTLFVDSYSE MLFFLQSLFM LATVVLYFSH LKEYVASMVF SLALGWTNML YYTRGFQQMG
rat TRPV1            KSLFVDSYSE ILFFVQSLFM LVSVVLYFSQ RKEYVASMVF SLAMGWTNML YYTRGFQQMG
dog TRPV1            KTLFVDSYSE MLFFVQSLFM LGTVVLYFCH HKEYVASMVF SLAMGWTNML YYTRGFQQMG
guinea pig TRPV1     KTLFVDSYSE ILFFVQSLFL LASVVLYFSH RKEYVACMVF SLALGWTNML YYTRGFQQMG
horse TRPV1          KTLFVDSYSE MLFFVQSLFM LGTVVLYFCH CKEYVASMVF SLAMGWTNML YYTRGFQQMG
human TRPV5          KTILGGPFHV IIIT-YASLV LVTMVMRLTN TNGEVVPMSF ALVLGWCSVM YFTRGFQMLG

                            K571 (rat)  R579 (rat)
                     601
human TRPV1          IYAVMIEKMI LRDLCRFMFV YIVFLFGFST AVVTLIEDGK NDSLPSESTS HRWRGPACRP
rat TRPV1            IYAVMIEKMI LRDLCRFMFV YLVFLFGFST AVVTLIEDGK NNSLPMESTP HKCRGSACKP
dog TRPV1            IYAVMIEKMI LRDLCRFMFV YLVFLFGFST AVVTLIEDGK NNSVPTESTL HRWRGPGCRP
guinea pig TRPV1     IYAVMIEKMI LRDLCRFMFV YLVFLFGFST AVVTLIEDGK NESLSAE--P HRWRGPGCRS
horse TRPV1          IYAVMIEKMI LRDLCRFMFV YLVFLFGFST AVVTLIEDGK NDSVLAESTS HRWRGHGCRS
human TRPV5          PFTIMIQKMI FGDLMRFCWL MAVVILGFAS AFYIIFQTED PTSLGQ---- ----------

                     661
human TRPV1          PDSSYNSLYS TCLELFKFTI GMGDLEFTEN YD--FKAVFI ILLLAYVILT YILLLNMLIA
rat TRPV1            -GNSYNSLYS TCLELFKFTI GMGDLEFTEN YD--FKAVFI ILLLAYVILT YILLLNMLIA
dog TRPV1            PDSSYNSLYS TCLELFKFTI GMGDLEFTEN YD--FKAVFI ILLLAYVILT YILLLNMLIA
guinea pig TRPV1     AKNSYNSLYS TCLELFKFTI GMGDLEFTEN YD--FKAVFI ILLLAYVILT YILLLNMLIA
horse TRPV1          PDSSYNSLYS TCLELFKFTI GMGDLEFTEN YD--FKAVFI ILLLAYVILT YILLLNMLIA
human TRPV5          -FYDYPMALF TTFELF---- -LTVIDAPAN YDVDLPFMFS IVNFAFTIIA TLLMLNLFIA

                            K688 (rat)
                     721
human TRPV1          LMGETVNKIA QESKNIWKLQ RAITILDTEK SFLKCMRKAF RSGKLLQVGY TPDGKDDYRW
rat TRPV1            LMGETVNKIA QESKNIWKLQ RAITILDTEK SFLKCMRKAF RSGKLLQVGF TPDGKDDYRW
dog TRPV1            LMGETVNKIA QESKNIWKLQ RAITILDTEK SFLKCMRKAF RSGKLLQVGY TPDGKDDYRW
guinea pig TRPV1     LMGETVNKIA QESKNIWKLQ RAITILDTEK SFLKCMRKAF RSGKLLQVGY TPDGKDDYRW
horse TRPV1          LMGETVNKIS QESKNIWKLQ RAITILDTEK SFLKCIRKAF RSGKLLQVGY TPDGKDDYRW
human TRPV5          MMGDTHWRVA QERDELWRAQ VVATTVMLER KLPRCLWPRS G---ICGCEF ----GLGDRW

                     781
human TRPV1          CFRVDEVNWT TWNTNV---G II-------- NEDP------ GNCEGVKRTL SFSLR-----
rat TRPV1            CFRVDEVNWT TWNTNV---G II-------- NEDP------ GNCEGVKRTL SFSLR-----
dog TRPV1            CFRVDEVNWT TWNTNV---G II-------- NEDP------ GNCEGIKRTL SFSLR-----
guinea pig TRPV1     CFRVDEVNWT TWNTNV---G II-------- NEDP------ GNCEGVKRTL SFSLR-----
horse TRPV1          CFRVDEVNWT TWNTNV---G II-------- NEDP------ GNCEDIKRTL SFSLR-----
human TRPV5          FLRVENHNDQ NPLRVLRYVE VFKNSDKEDD QEHPSEKQPS GAESGTLARA SLALPTSSLS

                     841
human TRPV1          ---SSRVSGR HWKNFALVPL LREASARDRQ SAQPEEVYLR QFSGSLKPED AEVFKSPAAS
rat TRPV1            ---SGRVSGR NWKNFALVPL LRDASTRDRH ATQQEEVQLK HYTGSLKPED AEVFKDSMVP
dog TRPV1            ---SGRVSGR NWKNFSLVPL LRDASTRERH PAQPEEVHLR HFAGSLKPED AEIFKDPVGL
guinea pig TRPV1     ---SGRVSGR NWKNFALVPL LRDASTRDRH SAQPEEVHLK HFSGSLKPED AEVFKDSAVP
horse TRPV1          ---SGRVSGR NWKNFSLAPL LREASVRERH PAPPEEVNLR HFAGSLKPED AEICKDPAAS
human TRPV5          RTASQSSSHR GWEILRQNTL GHLNLGLNLS EGDGEEV--Y HF-------- ----------

                     901
human TRPV1          GEK
rat TRPV1            GEK
dog TRPV1            GEK
guinea pig TRPV1     GEK
horse TRPV1          GEK
human TRPV5          ---
```

Figure S2

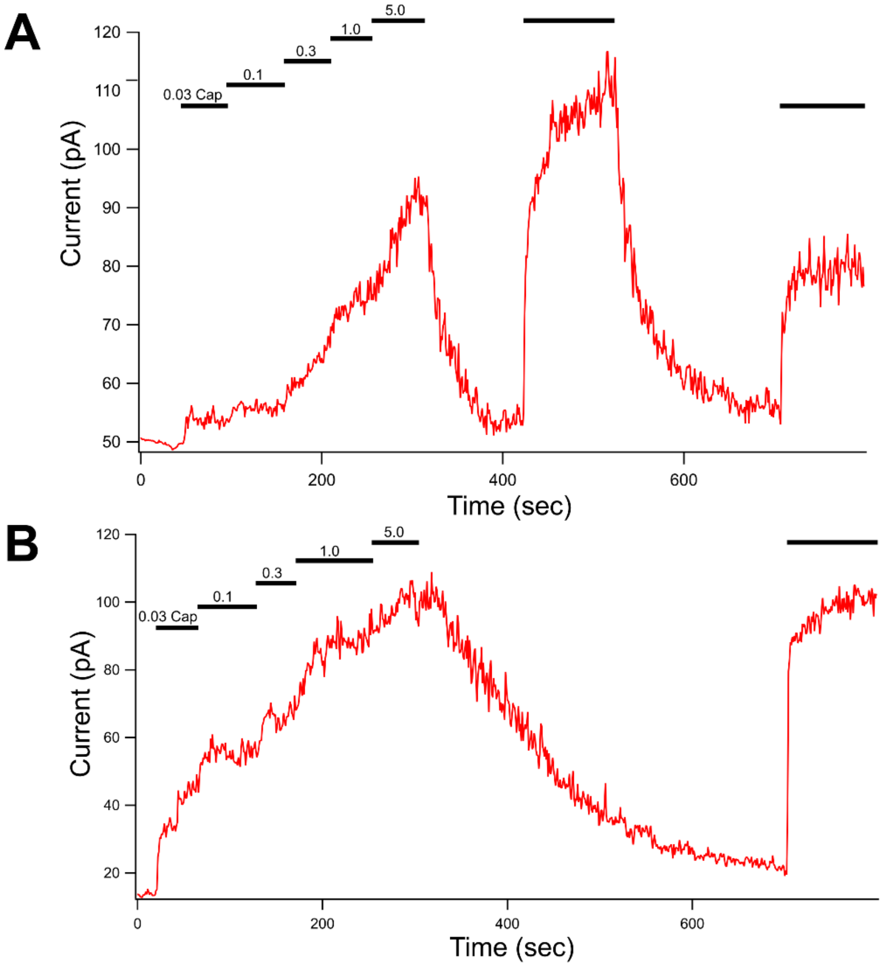
A
0.03 Cap
0.1
0.3
1.0
5.0
Current (pA)
Time (sec)
B
0.03 Cap
0.1
0.3
1.0
5.0
Current (pA)
Time (sec)

Figure S3

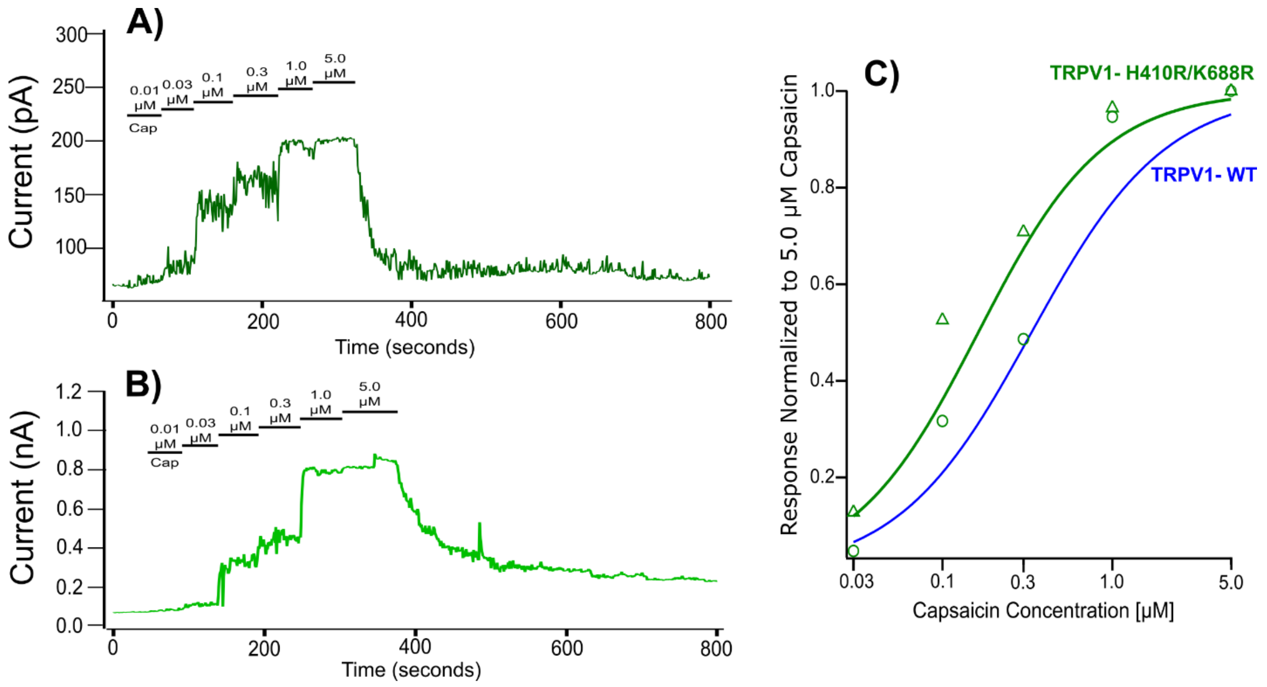
A)
Current (pA)
300
250
200
150
100
0.01 μM Cap
0.03 μM
0.1 μM
0.3 μM
1.0 μM
5.0 μM
0
200
400
600
800
Time (seconds)
B)
Current (nA)
1.2
1.0
0.8
0.6
0.4
0.2
0.0
0.01 μM Cap
0.03 μM
0.1 μM
0.3 μM
1.0 μM
5.0 μM
0
200
400
600
800
Time (seconds)
C)
Response Normalized to 5.0 μM Capsaicin
1.0
0.8
0.6
0.4
0.2
TRPV1- H410R/K688R
TRPV1- WT
0.03
0.1
0.3
1.0
5.0
Capsaicin Concentration [μM]

Table S1

| Residue (human numbering) | Gnomad 4.10 Variants (counts) |
|---|---|
| TRPV5-R305 | Histidine(21), Cysteine(9) |
| TRPV1-H411* | Glutamine(1), Arginine(1) |
| TRPV1-R410 | Histidine(294),Cysteine(12) |
| TRPV5-R492 | Histidine(25), Cysteine(18), Serine(1), Glycine(1) |
| TRPV1-R579 | Histidine(60), Cysteine(30), Leucine(2) |

*H411 in human is H410 in rat numbering
